# A Window into Mammalian Basement Membrane Development: Insights from the *mTurq2-Col4a1* Mouse Model

**DOI:** 10.1101/2023.09.27.559396

**Authors:** Rebecca A. Jones, Brandon Trejo, Parijat Sil, Katherine A. Little, H. Amalia Pasolli, Bradley Joyce, Eszter Posfai, Danelle Devenport

## Abstract

Basement membranes (BMs) are specialized sheets of extracellular matrix that underlie epithelial and endothelial tissues. BMs regulate traffic of cells and molecules between compartments, and participate in signaling, cell migration and organogenesis. The dynamics of mammalian BMs, however, are poorly understood, largely due to a lack of models in which core BM components are endogenously labelled. Here, we describe the *mTurquoise2-Col4a1* mouse, in which we fluorescently tag collagen IV, the main component of BMs. Using an innovative Planar-Sagittal live imaging technique to visualize the BM of developing skin, we directly observe BM deformation during hair follicle budding and basal progenitor cell divisions. The BM’s inherent pliability enables dividing cells to remain attached to and deform the BM, rather than lose adhesion as generally thought. Using FRAP, we show BM collagen IV is extremely stable, even during periods of rapid epidermal growth. These findings demonstrate the utility of the *mTurq2-Col4a1* mouse to shed new light on mammalian BM developmental dynamics.

## Introduction

The basement membrane (BM) is an ancient specialized extracellular matrix present in all metazoans (Özbek et al., 2010). It is a thin, dense sheet (∼50-100 nm) that underlies all epithelial and endothelial tissues. The BM serves multiple functions, including structural support, maintenance of tissue integrity and regulation of cell adhesion, migration, proliferation, and differentiation (Yurchenco, 2011). It is a complex structure composed of a network of proteins, glycoproteins, and proteoglycans and consists of three main layers: the lamina lucida, lamina densa (also called the basal lamina), and in some structures, the lamina fibro-reticularis (Timpl, 1996). The lamina lucida is in direct contact with the basal surface of epithelial or endothelial cells, and is made up primarily of laminins, nidogen and other glycoproteins, which are essential for cell adhesion and migration (Stanley et al., 1982, Schwarzbauer, 1999). The lamina densa, composed predominantly of type IV collagen, provides structural stability and acts as a barrier, separating the epithelial/endothelial compartment from the underlying mesenchyme (Pozzi et al., 2017).

Importantly, the BM acts as a selective filter, controlling the passage of cells and molecules between these different tissue compartments (Kelley et al., 2014). Many normal and pathological events involve cells transiting across basement membranes, including immune cell infiltration, cell migration during development, epithelial to mesenchymal transition and cancer metastasis (Morrissey and Sherwood, 2015, Nakaya and Sheng, 2013, Micalizzi et al., 2010, Ratzinger et al., 2002). As well as acting as a selective barrier, BMs are essential for cell migration, as they provide an active substrate to which cells can adhere. Laminins in BMs act as ligands to integrin receptors, coupling the extracellular environmental cues to the actin cytoskeleton of cells (Hynes, 1992, Sekiguchi and Yamada, 2018b). In doing so, the BM also regulates cell behaviors. In terms of cell guidance, the BM is a reservoir of signaling cues and matrix molecules that guide many cell types along their migration path, including neurons during axonal pathfinding and neural crest derived cells (Yasunaga et al., 2010, Leonard and Taneyhill, 2020, Banerjee et al., 2013). Tissue morphogenesis itself is regulated by BMs, via both molecular and biophysical processes. In the *Drosophila* embryo, knockdown of collagen IV reduces adult wing size due to a reduction in BMP signaling (Ma et al., 2017) and heterogeneity of BM stiffness in *Drosophila* is important for egg elongation (Chlasta et al., 2017). Overall, the BM performs a wide variety of functions including as a barrier, substrate, growth regulator and signaling platform, from early development throughout the whole adult life (Sherwood, 2021).

Studying BM dynamics or the crossing of cells through the BMs *in vivo* is particularly challenging. BM crossing events are stochastic in nature and occur deep within tissues. Therefore, much of what is known about how cells cross BMs is derived from *in vitro* studies (Glentis et al., 2017, Bahr et al., 2022, Albini and Noonan, 2010), which, although informative, still require validation in the *in vivo* context. Our knowledge of the *in vivo* dynamics of BMs has come from elegant experiments in *Drosophila* and *C. elegans*, in which endogenous components of the BM have been fluorescently labelled. The *Vkg::GFP* (collagen IVa2) *Drosophila* line, created as part of a protein-trap strategy (Morin et al., 2001, Buszczak et al., 2007), has been invaluable in studying dissemination of tumor cells through the BM, as well as in BM-controlled shaping of growing organs during development (Tamori et al., 2016, Rei et al., 2021, Harmansa et al., 2023, Page-McCaw et al., 2007, Page-McCaw, 2008). The stereotyped invasion of the *C. elegans* anchor cell through the uterine and vulval basement membranes during development, together with the successful endogenous tagging of over 60 *C. elegans* BM proteins has greatly increased our understanding of BM composition, dynamics and the cellular and molecular mechanisms underpinning the invasion process (Schindler and Sherwood, 2013, Sherwood and Sternberg, 2003, Costa et al., 2023, Kelley et al., 2017, Keeley et al., 2020, Kelley et al., 2019, Kenny-Ganzert and Sherwood, 2023).

A key bottleneck in our understanding of BM dynamics in mammals (and indeed vertebrates) is the inability to visualize the BM live, *in vivo*. Tagging endogenous BM components presents unique challenges because many are subject to extensive proteolytic cleavage and post-translational modifications (Frantz et al., 2010). Moreover, the addition of a fluorescent tag to a protein monomer that is subsequently highly crosslinked and integrated into a complex and dense protein network can interfere with the formation of that network via steric hindrance. Several attempts to visualize the vertebrate BM *in vivo* have been made, for example the *lamC1:lamC1-sfGFP* zebrafish line recapitulates the reported mRNA expression pattern of endogenous *lamC1,* but the tag itself is not inserted into the endogenous genomic locus and the fusion protein is expressed as an additional copy (Yamaguchi et al., 2022, Sztal et al., 2011). Similarly, Futaki et al., (2023) have generated a R26-CAG-Nid1-mCherry mouse line in which the BM can be visualized in *vivo*, however again this tag is not inserted in the endogenous locus; the fusion protein is expressed from a ubiquitous promoter and the model shows ectopic expression of Nid1-mCherry as well as reduced localization of the endogenous Nid1. More recently, Morgner et al., (2023) generated an endogenous *Lamb1-Dendra2* mouse model in which heterozygous animals have fluorescently labelled laminin β-1-containing BMs, which enabled tumor-cell mediated BM turnover to be measured using intravital microscopy. A limitation of this model, however, is that only BMs containing the laminin β-1 isoform can be visualized. In order to visualize all BMs and truly appreciate the structural complexity of the BM as a barrier, the ideal component to fluorescently tag is collagen IV, as it makes up ∼50% of all BMs (LeBleu et al., 2007).

Collagen IV is a heterotrimeric molecule, expressed exclusively in BMs, with all the collagen IV in mammals being derived from six different α-chains, encoded by the genes *COL4A1*, *COL4A2*, *COL4A3*, *COL4A4*, *COL4A5* and *COL4A6* (Netzer et al., 1998, Mak and Mei, 2017). All BMs are composed solely of α1 and α2 chains during development, with some specialized structures adding a secondary network during maturation, such as α3/α4/α5 in the kidney glomerulus (Kruegel and Miosge, 2010). Collagen IV is not required to initiate BM deposition during development, but loss of both *Col4a1* and *Col4a2* is embryonically lethal in mice by E10.5, due to lack of BM structural integrity and the breakdown of Reichert’s membrane (Pöschl et al., 2004). As the α1 and α2 chains are ubiquitously expressed in BMs, we sought to label the endogenous *Col4a1* gene locus in mouse, to enable all BMs to be visualized.

In this work, we present the novel *mTurquoise2-Col4a1* (*mTurq2-Col4a1*) mouse line, in which endogenous collagen 4a1 (COL4A1), and therefore all BMs, are labelled. Despite being homozygous lethal, which is a developmental phenotype that requires further analysis, heterozygous animals are healthy and fertile, with normal BM architecture and composition in the dermal-epidermal junction BM of the skin, suggesting incorporation of one copy of the mTurquoise2 protein at the endogenous locus of *Col4a1* can support normal functionality of murine BMs. Furthermore, the stability and brightness of mTurquoise2 means BMs are clearly visible in heterozygous animals, allowing for long-term live imaging to be undertaken. Using an innovative mounting and imaging methodology, we show that the dermal-epidermal junction BM (hereafter referred to as the epidermal BM for simplicity) in embryonic mouse skin can be imaged long-term using explants from *mTurq2-Col4a1* mice. This has revealed new insights into epidermal BM morphology and behavior during hair follicle development suggesting a previously underappreciated inherent pliability. Moreover, this flexibility of the epidermal BM allows dividing basal progenitor cells to maintain adhesion to the BM even during division, challenging the notion that all dividing cells reduce or lose their adhesion to the substrate when they round up in metaphase. We describe novel observations of COL4A1 in the dermis, including a potential role in stabilizing the developing dermal condensate. Finally, we show that despite the significant growth that occurs during E13.5 and E15.5, COL4A1 is strikingly stable in the BM of the developing epidermis, suggesting the inherent pliability observed may be what allows for growth and morphogenetic changes, rather than extensive proteolysis and degradation of the core components. We envisage that the *mTurq2-Col4a1* mouse model will be a useful addition to the field of BM research as it allows direct visualization of BM assembly, deformation and remodeling during development, and should be applicable to pathological conditions. Moreover, this model will allow many of the important observations made *in vitro* and in invertebrates to be visualized directly in the clinically relevant mouse model.

## Material and Methods

### Generation and breeding of the *mTurq2-Col4a1* transgenic mouse line

The *mTurq2-Col4a1* mouse line was generated by knocking-in the coding sequence of *mTurquoise2* after the START codon and signal sequence of the endogenous *Col4a1* genomic locus to create an N-terminal fusion reporter using the previously described 2C-HR CRISPR method (Gu et al., 2020, Gu et al., 2018). Briefly, an HR repair plasmid was constructed by amplifying homology arms 1000 bp upstream and downstream of the mTurquoise2 insertion site by PCR from genomic DNA obtained from murine CD1 keratinocytes and inserted using InFusion cloning (Takara) into the mTurquoise2-N1 plasmid (Addgene plasmid #54843;http://n2t.net/addgene:54843;RRID:Addgene_54843, a gift from Michael Davidson and Dorus Gadella). Insertion was designed to remove the STOP codon at the 3’ end of the mTurquoise2 ORF. Site directed mutagenesis was then performed to silently mutate the PAM sites using NEB Q5 Site-Directed Mutagenesis kit (E0554) according to the manufacturer’s instructions. sgRNAs were designed using CRISPOR (Concordet and Haeussler, 2018) and two were selected based on high predicted cutting efficiencies (sgRNA1 and sgRNA2). Synthetic sgRNAs were then ordered from Synthego with chemical modifications at the 5′ and 3′ termini: 2′-O-methyl modified bases and 3′ phosphorothioate linkages; for sequences see table S1.

CD1-IGS mice (Charles River strain 022) were used as embryo donors. Briefly, female CD1-IGS were super-ovulated at 5-7 weeks of age using 7.5 IU PMSG (Biovendor) administered by IP injection followed by 7.5 IU HCG (Sigma) by IP injection 47 hours post PMSG. Super-ovulated females were mated to CD1-IGS stud males and checked for copulatory plugs the following morning. Cytoplasmic microinjection of 2-cell embryos was performed as previously described (Gu et al., 2020, Gu et al., 2018). Briefly, embryos were harvested at the 2-cell stage at E1.5 by flushing the oviducts with M2 Media (Cytospring) and each cell was microinjected with 100 ng/ul *Cas9* mRNA (made by *in vitro* transcription (mMESSAGE mMACHINE SP6 transcription kit, Thermo Fisher) using Addgene plasmid 122948), 30 ng/ul donor plasmid and 50 ng/ul sgRNA, using a Leica Dmi8 inverted epifluorescent microscope, an Eppendorf Femtojet and a Micro-ePore (WPI). Embryos were immediately transferred into the oviducts of pseudo-pregnant female CD1 mice. Positive founders (N_0_) were identified by PCR genotyping as follows. gDNA from pup ear punches was prepared using Purelink Genomic DNA Mini Kit (Invitrogen), before being used as template in two PCR reactions. Genotyping PCRs were designed to i) only amplify a fragment if insertion had occurred at the correct locus and ii) differentiate between hetero-and homozygotes by spanning the entire insertion area. See table S1 for primer sequences. N_0_ founders were then crossed to C57BL/6J mice, and N_1_ mice from these crosses were genotyped as above using a proof-reading polymerase (NEB Q5 DNA Polymerase). Purified amplicons from these PCR reactions were then Sanger sequenced to ensure accuracy of insertion site and genomic sequence. N_1_ *mTurq2-Col4a1* heterozygotes were then outcrossed to C57BL/6J mice for a minimum of five generations. Heterozygote incrosses from all generations did not yield any homozygous offspring. Mice were housed in an AAALAC-accredited facility following the Guide for the Care and Use of Laboratory Animals. Animal maintenance and husbandry followed the laboratory Animal Welfare Act. Princeton University’s Institutional Animal Care and Use Committee (IACUC) approved all animal procedures and the 3 ‘R’s were considered in all experimental design.

### Immunofluorescence and fixed sample imaging

The following mouse tissues were embedded fresh in OCT on dry ice and stored at −80°C: E18.5 embryos, dissected in PBS and heads removed; adult kidneys, whole; adult back skin, shaved using clippers to remove the hair and dissected into strips along the midline. Frozen OCT-embedded tissues were cryosectioned on a Leica CM3050 cryostat. Sagittal sections were cut at 10 µm thickness, collected on Superfrost Plus slides and allowed to dry before being stored at −20°C. For immunostaining, sections were moved to a slide holder container and fixed in 4% paraformaldehyde (PFA) in PBS for 10 mins at room temperature (RT), washed in PBS for 5 mins, permeabilized in PBT2 (PBS with 0.2% Triton X-100) for 10 mins at room temperature (RT), and blocked in 2% normal goat serum, 2% normal donkey serum and 2% bovine serum albumin (BSA) in PBT2 overnight at 4°C with gentle rocking. Slides were moved to a humidified chamber on a flat surface and incubated with primary antibody in block for 1 hour at RT; see table S2 for primary antibodies and concentrations used. Slides were washed in slide holder containers three times for 10 mins with PBT2 and incubated in humidified chamber on a flat surface with secondary antibodies in PBT2 for 1 hour at RT. Alexa Fluor-555 and −647 secondary antibodies were used at 1:2000 (Invitrogen). Slides were washed in slide holder containers in PBT2 with Hoechst (1 µg/mL, Invitrogen, H1399) for 10 mins, followed by two 10 mins washes in PBT2. A final wash in PBS for 5 mins was carried out before mounting in ProLong Gold (Invitrogen, P36930). Slides were stored at −20°C until imaged.

For immunofluorescence of E15.5 and E12.5 backskins, embryos were dissected in PBS (+Mg,+Ca) and fixed in 4% PFA for 1 hour at RT with gentle rocking. Backskins were then incubated in blocking medium overnight at 4°C with gentle rocking (1% BSA, 1% fish gelatin, 2% normal goat serum and 2% normal donkey serum in PBT2 with 0.01% sodium azide). The following day, samples were incubated in primary antibody (in blocking medium) overnight at 4°C with gentle rocking. See table S2 for primary antibody details and concentrations used. Samples were then washed a minimum of five times (30 mins per wash) in PBT2 before incubating overnight in secondary antibodies (Alexa Fluor-555 and −647, 1:1000, Invitrogen) in PT2 at 4°C with gentle rocking in the dark. Following secondary incubation, samples were washed a minimum of five times in PBT2 (30 mins per wash) with the first wash containing Hoeschst (Invitrogen, H1399, 1 µg/ml), before being mounted in ProLong Gold (Invitrogen, P36930).

Images were acquired on an inverted Nikon A1R confocal microscope controlled by NIS Elements software. Objectives used were Plan Apo 10/0.45NA air, Plan Apo 20/0.75NA air and Plan Apo 60/1.4NA oil immersion (Nikon). For XZ optical reconstructions, resonance imaging was used with 8x averaging and images denoised in NIS Elements. NIS Elements and FIJI were used for all image processing.

### Pearson’s Correlation Coefficient and overlap fraction analyses

Images were acquired as described above using a Plan Apo 60/1.4NA oil immersion objective with 6x zoom to obtain a pixel size of 0.09 µm and an optical resolution of 0.26 µm (to attain Nyquist). Z stacks were obtained with 0.3 µm step size over a range of 3 µm with the center of the stack being set at the epidermal BM. 10 individual z stacks were obtained per embryo (n=3). To calculate Pearson’s Correlation Coefficient, the Z slice at which the epidermal BM was most in focus was selected, background subtracted and the FIJI plugin Coloc 2 (Costes threshold regression; Costes randomisations = 100) was used on a region of interest (ROI) around the area of laminin β-1 signal (avoiding areas devoid of signal to reduce noise). Both biological (n=3) and technical (n=30) replicates were plotted and statistical analysis undertaken using GraphPad Prism.

Overlap fraction analysis was undertaken using a custom macro in FIJI. Single Z planes in both channels as described above were thresholded using the Otsu method to generate a mask. The overlap map was then thresholded (Otsu) and an ROI generated around the entire overlap region. The area of the overlap region was then divided by the area of the mask for each channel to calculate the overlap fraction. Both biological (n=3) and technical (n=30) replicates were plotted and statistical analysis undertaken using GraphPad Prism.

### Transmission Electron Microscopy (TEM)

E18.5 embryos were harvested from the euthanized dam and back skins immediately dissected into fresh TEM fixative (2% glutaraldehyde, 4% formaldehyde, 2mM CaCl_2_ in 0.1 M sodium cacodylate buffer, pH 7.2). Samples were gently rocked at RT for 1 hour, before storage at 4°C until processed for imaging. Samples were postfixed in 1% osmium tetroxide and processed for Epon embedding. Ultrathin sections (60–65 nm) were counterstained with uranyl acetate and lead citrate. Images were taken with a transmission electron microscope (Tecnai G2-12; FEI, Hillsboro, OR) equipped with a digital camera (AMT BioSprint29).

### Planar-Sagittal (PS) Multiview live imaging

A stencil to create a ridge in the agarose gel was created using Conair^TM^ bobby pins and a standard glass microscope slide (Fisherbrand). Three bobby pins were cut at the apex, then the flat arms from each pin were super glued side-on, to a microscope slide. This created a flat surface with a large bar in the center measuring 3 mm wide, 25 mm long and 1 mm tall. F-medium agarose plates were then made as previously described (Cetera et al., 2018) with the following additional steps. After briefly cooling to 60 °C, the liquid agarose-medium was poured into a 35 mm dish plate at 4°C. After 2 mins had elapsed, the bobby pin glass stencil was dropped onto the solidifying gel, allowing it to float on top. After 15 mins at RT, the bobby pin glass stencil was carefully removed from the gel using forceps. The ridged gel was then stored in a 37°C, 5% CO_2_ cell-culture incubator while dissections took place. E14.5 and E15.5 flank skin explants were dissected in PBS (+Mg,+Ca) as previously described (Cetera et al., 2018) and transferred to the agarose-F-medium gel with the ridge. The explants were mounted dermal side down with approximately 50% of the lateral flank region being placed over the edge of the ridge, and the midline region mounted firmly on the flat portion of the gel to prevent slipping into the ridge. Explant gels were then mounted into Lumox membrane dishes (Sarstedt) as previously described. Z-stacks with 3 µm step sizes were then acquired over several XY positions at 7-min intervals for 5-15 hours using a Nikon inverted CSU-W1 SoRa Spinning Disk microscope with a Plan Apo Lambda S 40XC Sil/1.25NA immersion objective, controlled by NIS Elements software. Sample temperature was maintained during imaging using a Tokai Hit humidified stage top incubator and an environmental imaging chamber, both set to 37°C with 5% CO_2_. Throughout all imaging experiments, areas that were close to the edge of the explants were not imaged to avoid any wound healing response. NIS Elements and FIJI were used for movie processing.

### Live-image analysis and quantification

The mTurq2-COL4A1 signal intensity was determined during PS Multiview imaging using FIJI software. An ROI was drawn both around the mTurq2-COL4A1 signal at the base of the hair follicle and around the interfollicular epidermis (IFE) region in view. If the hair follicle was in the center of the image, two IFE ROIs were drawn on either side of the hair follicle and the mean values obtained. The average intensity of the mTurq2-COL4A1 signal in both hair follicle and IFE ROIs was then measured, and the ratio between the two was then calculated for each time point across the different movies (n=8 movies). The two stages of hair follicle development (placode and hair germ) were determined by measuring the depth of the hair follicle relative to the IFE at every time point across all movies, then binning the different depths into two equal groups, representing the two developmental stages. The background IFE/IFE ratio was determined by taking each individual IFE mTurq2-COL4A1 signal intensity measurement and dividing it by the average IFE mTurq2-COL4A1 signal for the corresponding movie. Graphing and statistical analysis was undertaken using GraphPad Prism.

To measure BM deformation occurring during cell division, we identified dividing basal progenitor cells and measured the angle between the dividing cell’s most basal contact with that of both of its closest neighbors. The center of the basal surface of the dividing cell and the two neighboring cells, together with the basal most intersection of the two neighboring cells, were used as the points from which to measure the angles. The angles were measured at three time points: 18 mins before the onset of division when the cell is in interphase (columnar), during metaphase when the cell rounds up, and 18 mins after the daughter cells separate. The mean of both angles at each time point was calculated and plotted, n=23 dividing cells that were followed through division. Graphing and statistical analysis was undertaken using GraphPad Prism.

### Fluorescence recovery after photobleaching (FRAP)

BMs in live skin explants from E13.5 or E15.5 *mTurq2-Col4a1/+* embryos were imaged using a Plan Apo 20/0.45NA air objective (Nikon) with appropriate optical zoom on an inverted Nikon A1R laser scanning confocal microscope equipped with a stage-top Tokai Hit incubation chamber to maintain 37°C and 5% CO_2_. 1.5 µm diameter circular ROIs were bleached using a 445 nm laser and recovery was monitored for a period of 10 mins with 10-second intervals from either IFE or along the rim of developing hair follicles. Three reference pre-bleach images were also acquired. Magnification, laser power (both for bleach and acquisition), pixel dwell time and acquisition rate were kept uniform across all measurements. The acquired images and measured ROIs in the time series were checked for Z-drift and corrected for presence of any XY drift. A reference ROI was made in a non-bleached region to correct for overall bleaching during image acquisition. Background autofluorescence was measured from non-fluorescent wild type skin using the same image acquisition conditions. The ROI values were extracted from drift corrected images in NIS Elements software and subsequently processed in Microsoft Excel and GraphPad Prism. Each image time series was background/autofluorescence subtracted and bleach corrected (to be referred to as corrected intensity henceforth) and thereafter the corrected intensity profile was normalized as (F_t_ – F_bleach_)/ (F_ini_ – F_bleach_), where, F_t_ is the corrected intensity of the ROI at a given time point, F_bleach_ is the corrected intensity at the time point immediately after bleaching and F_ini_ is the mean ROI intensity of the three pre-bleach frames. Each mean recovery curve was fitted to exponential one phase association equation in GraphPad Prism with an r-squared value > 0.9 to determine the fitted plateau and Y_0_ values, which were then used to determine the immobile fraction = 1-[(Plateau-Y_0_)/(1-Y_0_)]. Data represented is pooled from three biological replicates.

### Supplemental material

Figure S1, related to Figure 1, shows an additional experimental replicate of mTurq2-COL4A1 labelling in adult mouse skin and kidney. Figure S2, related to Figure 3, shows TEM images from additional experimental replicates of E18.5 WT and *mTurq2-Col4a1/+* embryonic skin. Figure S3, related to Figure 5, shows additional examples of snapshots taken from time-lapse imaging of BM curvature around dividing basal progenitor cells in *mTurq2-Col4a1/+;mTmG*/*mTmG* backskins, and quantification of average angle of deformation. Movie S1 shows three representative time-lapse live-imaging movies of hair follicle invagination in E14.5-E15.5 *mTurq2-Col4a1/+;mTmG*/*mTmG* backskins. Movie S2 shows three representative time-lapse live-imaging movies of basal progenitor cell divisions in E14.5-E15.5 *mTurq2-Col4a1/+;mTmG*/*mTmG* backskins. Movie S3 shows a Z-stack movie in planar view through a fixed E15.5 *mTurq2-Col4a1/+* backskin from the basal layer (basement membrane) through to the dermis. Table S1 details sgRNA and oligonucleotide sequences used in this study; Table S2 details primary antibodies used.

## Results

### Generation of the *mTurquoise2-Col4a1* mouse model

The collagen IV α chains are characterized by a central triple helical collagenous region of Gly-XY repeats, flanked by a 7S domain at the amino terminal and a globular NC1 carboxy domain (Figure 1A). The central collagenous region confers both structural stability and flexibility to the molecule, the NC1 domain is required to assemble the collagen α1α1α2(IV) heterotrimer, and the 7S domain is involved in the assembly of the formed heterotrimers into a more complex lattice network to form the BM (Timpl, 1989, Mats Paulsson, 2008). It therefore follows that inserting a fluorescent protein in any position that interferes with or disrupts these domains may interfere with protein function and/or proper assembly of the BM. To inform our strategy, we studied the position of the *Drosophila Vkg::GFP* protein trap insertion, which is viable and labels all BMs (Morin et al., 2001, Buszczak et al., 2007). In this line, GFP is inserted between the N’ terminal signal sequence that is cleaved within the cell before secretion, and the highly conserved 7S region. We mapped the *Vkg::GFP* insertion locus to the analogous position in the mouse *Col4a1* gene, which corresponds to a position towards the 5’ end of exon 2 (Figure 1B), and a target insertion site on the COL4A1 protein between amino acids K28-G29.

**Figure 1.**
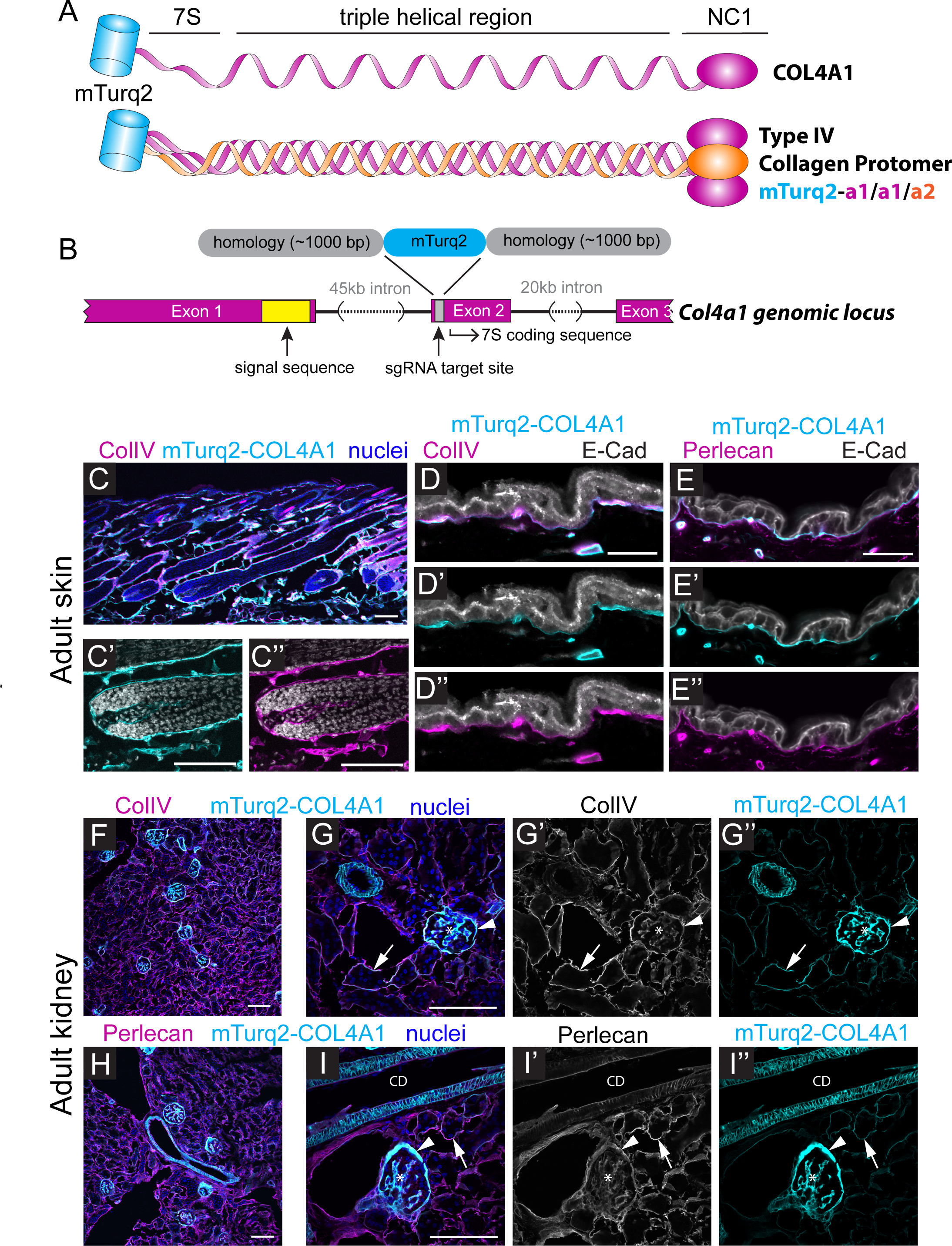
*mTurq2-Col4a1* endogenously tagged reporter design and basement membrane localization in adult mouse tissues. **(A)** Schematic of the COL4A1 subunit and type IV collagen trimeric protomer with an N-terminal mTurq2 fluorescent tag. **(B)** Schematic of the *Mus musculus Col4a1* genomic locus and *mTurq2* insertion site. The *mTurq2* gene was inserted into Exon 2 between the coding sequences for the signal peptide and 7S domain. **(C)** Sagittal section of dorsal skin from *mTurq2-Col4a1/+* adult mouse labelled with ColIV antibodies (magenta) and Hoechst to mark nuclei (blue; grayscale in C’-C’’). (C’) mTurq2-COL4A1 (cyan) localizes to BM surrounding the hair follicle, where it overlaps with ColIV staining (C’’, magenta). Scale bars 100 µm. **(D-D’’)** High magnification view of mTurq2-COL4A1 (cyan) localization in the interfollicular epidermis from *mTurq2-Col4a1/+* adult mouse labeled with ColIV (magenta) and E-Cadherin (E-Cad; grayscale). Scale bar 20 µm. **(E-E’’)** High magnification view of mTurq2-COL4A1 (cyan) localization in the interfollicular epidermis from *mTurq2-Col4a1/+* adult mouse labeled with Perlecan (magenta) and E-Cadherin (E-Cad; grayscale). Scale bar 20 µm. **(F)** Kidney section from *mTurq2-Col4a1/+* adult mouse labelled with ColIV (magenta) and Hoechst to mark nuclei (blue). Scale bar 100 µm. **(G-G’’)** High magnification view of mTurq2-COL4A1 (cyan) localization in the kidney of *mTurq2-Col4a1/+* adult mouse labeled with ColIV (magenta in G; grayscale in G’) Hoechst (blue). mTurq2-COL4A1 expression can be seen in renal tubules (arrows), Bowman’s capsule (arrowheads) and the mesangial matrix (asterisks). Scale bar 100 µm. **(H)** Kidney section from *mTurq2-Col4a1/+* adult mouse labeled with Perlecan (magenta) and Hoechst (blue). Scale bar 100 µm. **(I-I’’)** High magnification view of mTurq2-COL4A1 (cyan) localization in the kidney of *mTurq2-Col4a1/+* adult mouse labeled with Perlecan (magenta in G; grayscale in G’) and Hoechst (blue). mTurq2-COL4A1 expression can be seen in renal tubules (arrows), Bowman’s capsule (arrowheads), the mesangial matrix (asterisks) and the collecting duct (CD). Scale bar 100 µm.

Many fluorescent proteins, despite being termed monomeric, have a tendency to oligomerize under physiological conditions (Costantini et al., 2012). We therefore chose mTurquoise2 as our fluorescent protein tag, as it does not homodimerize under any of the currently established gold-standard tests (Goedhart et al., 2012). mTurquoise2 also has an ‘in-built’ linker region precluding the need for artificial linker sequences. Moreover, mTurquoise2 has a low acid sensitivity (pKa 3.1), meaning it is stable in the lower pH of the lumenal and extracellular space (Denda et al., 2000, Chan and Mauro, 2011). It is also bright and photostable (t½ = 90s compared with ∼50s for EGFP) and its excitation/emission spectrum means it can be used in combination with YFP, mCherry/tdTomato and other far-red fluorescent proteins (Lambert, 2019).

To insert the mTurquoise2 ORF into the target locus of the *Col4a1* mouse gene, we used highly efficient 2-cell homologous recombination CRISPR/Cas9 mediated mutagenesis as previously described (Gu et al., 2020, Gu et al., 2018). A circular plasmid repair template containing the mTurquoise2 ORF flanked by ∼1000 bp homology arms, together with a sgRNA and *Cas9* mRNA were injected into the cytoplasm of both blastomeres of 2-cell mouse embryos. Using this approach, we successfully generated an mTurquoise2-collagen 4a1 mouse line (*mTurq2-Col4a1*), in which the mTurquoise2 ORF is inserted downstream of the cleaved signal sequence and upstream of the conserved 7S domain (Figure 1B). Histological analysis of adult skin from *mTurq2-Col4a1/+* mice confirmed robust and clear localization of mTurq2-COL4A1 signal in the epidermal BM (Figure 1C-E and S1A). In the kidney, mTurq2-COL4A1 signal was observed in Bowman’s capsule, the mesangial matrix and tubular BMs. (Figure 1F-I and S1B). The mTurq2-COL4A1 signal in each tissue recapitulated the expression pattern of collagen IV antibody staining (Figure 1C”,D”,G’). We also labelled both these tissues for perlecan, the most abundant heparan sulfate proteoglycan in the BM (Figure 1E”,H,I’ and S1A-B). Aggregated perlecan connects the laminin polymers of the BM into the collagen IV network, in a ‘spot-welding’ manner, thus connecting these two distinct and critical BM networks (Behrens et al., 2012). The pattern of mTurq2-COL4A1 expression coincided with perlecan expression in all BMs observed.

### Characterization of mTurq2-COL4A1 expression in E18.5 mouse skin

The BM is established during embryonic development, with laminins expressed around gastrulation followed by collagen IV shortly thereafter (Yurchenco et al., 2004). Studies have shown that although collagen IV is not required to initiate BM assembly, it is required for functional stabilization of BMs from as early as E10.5 (Guo et al., 1991, Guo and Kramer, 1989, Pöschl et al., 2004). Moreover, unlike laminins which express additional subunit isoforms in post-natal life, collagen α1 and α2 chains are laid down mainly during embryogenesis, with only a small number of BMs replacing some of their α1 and α2 chains with α3/α4/α5 chains (Gunwar et al., 1998). We therefore examined the epidermal BM in embryonic mouse skin. Both WT and *mTurq2-Col4a1/+* embryos were labelled with antibodies against collagen IV (ColIV), laminin β-1 (Lamb1) and perlecan (Figure 2). The epidermal BM was clearly labelled in *mTurq2-Col4a1/+* embryonic skin (Figure 1B’’’ and 1E’’’), and its localization overlapped with that of the ColIV antibody (Figure 1B’’).

**Figure 2.**
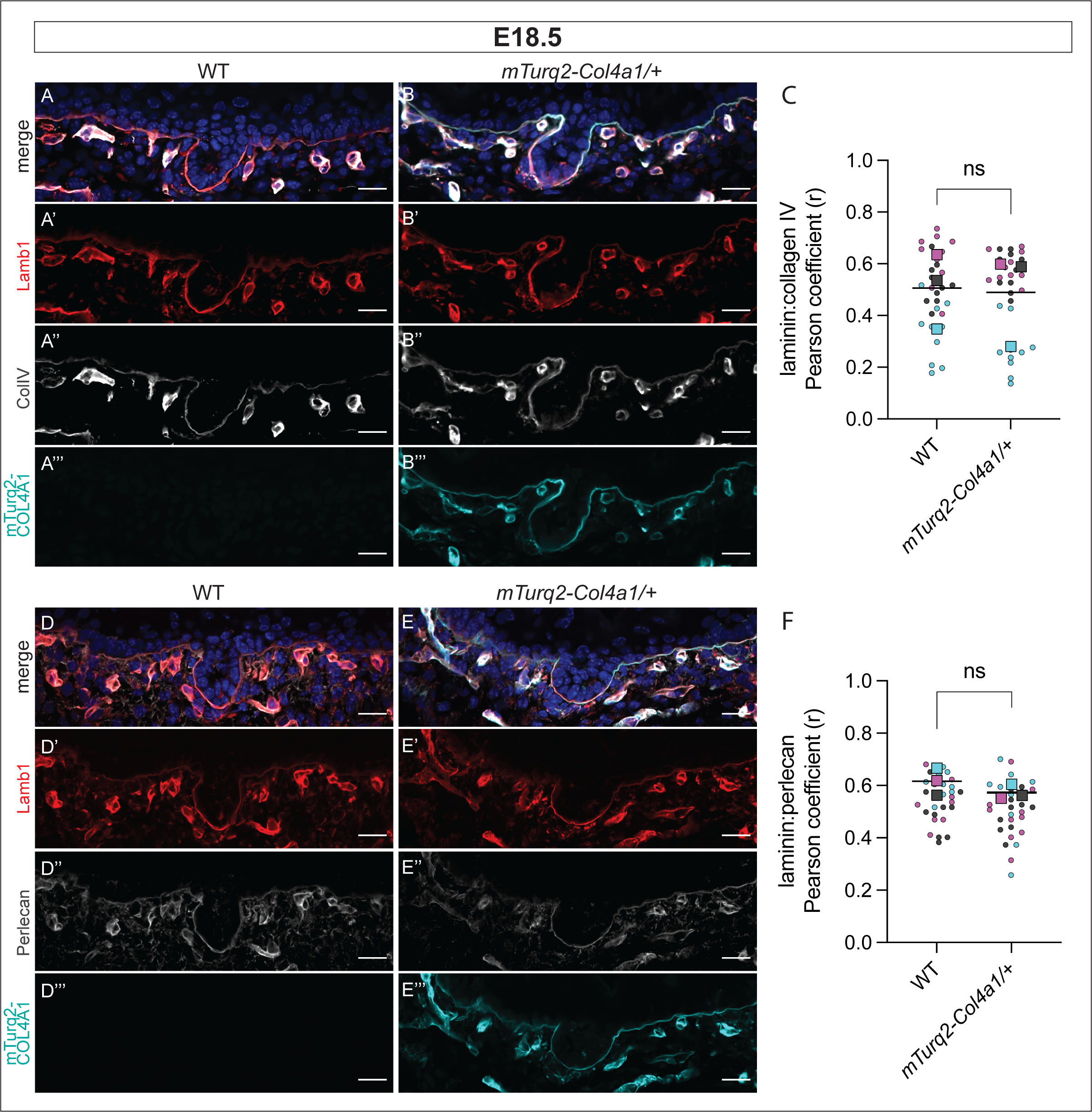
E18.5 *mTurq2-Col4a1* mouse backskin shows correct and robust expression of mTurq2-COL4A1 in the dermal:epidermal junction basement membrane. **(A)** Average intensity projection of backskin from WT E18.5 embryo labelled with Hoechst (merge, top), anti-laminin β-1 antibody (A’) and anti-collagen IV antibody (A’’). No signal is observed in the mTurq2 channel (A’’’). **(B)** As (A) except backskin from an *mTurq2-Col4a1/+* embryo. Robust mTurquoise2 signal recapitulates the labelling pattern of the collagen IV antibody. **(C)** Pearson’s correlation coefficient (r) measuring the correlation between laminin β-1 and collagen IV expression (antibody labelling) in WT and *mTurq2-Col4a1/+* E18.5 backskins as labelled. Larger datapoints – biological replicates (n=3 embryos), smaller datapoints – individual images per replicate (n=10 per embryo). **(D)** Average intensity projection of backskin from WT E18.5 embryo labelled with Hoechst (merge, top), anti-laminin β-1 antibody (D’) and anti-perlecan antibody (D’’). **(E)** As (A) except backskin from an *mTurq2-Col4a1/+* embryo. **(F)** As (C) except measuring the correlation between laminin β-1 and perlecan. Scale bar 20 µm. ns = not significant, Mann-Whitney U-test.

The spatial relationship between the two major structural components of the BM, the lamina lucida and the lamina densa, is critical to BM function. The supramolecular laminin containing network of the lamina lucida is linked to the collagen IV lamina densa network by the HSPG perlecan, with a potential role also played by agrin (Behrens et al., 2012, Hohenester and Yurchenco, 2013). In keeping with this, mutations in human *COL4A2* result in a failure of COL4A2 and laminin colocalization in the epidermal BM (Murray et al., 2014). Similarly, a 40% reduction in perlecan staining in all major BMs was observed in mice carrying *Col4a1* mutations (Jones et al., 2016). We therefore asked whether the relationships between these critical BM proteins are affected by the incorporation of mTurq2-COL4A1. To do so, we measured the linear correlation between Lamb1 and ColIV antibody labelling in WT and *mTurq2-Col4a1/+* E18.5 epidermal BMs at pixel-level scale in high resolution images (see Materials and Methods). As expected, we saw that the Pearson’s Correlation Coefficient (PCC) r value was variable, as both supramolecular networks will not occupy the same space, but crucially, no difference was detected between WT and *mTurq2-Col4a1/+* backskins, suggesting the *mTurq2-Col4a1/+* allele does not interfere with the network interactions of laminin β-1 and collagen IV (Figure 2C). We repeated this analysis measuring the correlation between Lamb1 and perlecan antibody labelling and observed that the perlecan localization profile in *mTurq2-Col4a1/+* E18.5 epidermal BMs resembled that of WT littermates (Figure 2D’’ and E’’, F). Taken together, these data demonstrate that the structure of the epidermal BM in *mTurq-Col4a1/+* embryos resembles that of WT littermates.

Despite numerous (>10) heterozygous crosses at several stages during development of the *mTurq2-Col4a1* line, we never observed the birth of any homozygous pups. This suggests that the *mTurq2-Col4a1* line is embryonically homozygous lethal, although we saw no apparent defects in heterozygous mice, which resemble WT littermates in terms of longevity, fertility, and overall health. The fact that heterozygous animals were viable, healthy and fertile suggested that incorporation of mTurq2-COL4A1 was not overtly deleterious to BM function when present as a single genomic copy. Furthermore, our correlation coefficient data in skin showed no difference in the spatial relationships between core BM components, as would be expected if incorporation of the mTurq2-COL4A1 protomer was detrimental to BM formation. However, despite attaining high resolution images to complete this analysis, our data were inherently limited by the resolution limits of light microscopy. To study the mTurq2-COL4A1 BM at the ultrastructural level therefore, we next turned to electron microscopy.

### Basement membranes in *mTurq2-Col4a1/+* mice resemble those of WT littermates at the ultrastructural level

Using transmission electron microscopy (TEM) we performed a detailed analysis of skin architecture and the BM at the dermal-epidermal junction of E18.5 *mTurq2-Col4a1/+* embryos (Figure 3). The overall skin architecture in *mTurq2-Col4a1/+* embryos was similar to WT littermates, consisting of five distinct layers (dermis, basal, spinous, granular and stratum corneum) that displayed characteristic morphologies associated with each layer (i.e. dense keratin bundles in the spinous layer, and keratohyalin granules in the granular layer) (Figure 3A-B, Figure S2). High magnification views of the junction between the basal epithelial layer and the underlying dermis showed numerous and regularly spaced BM thickenings at sites of hemidesmosome attachments (Figure 3C-F, Figure S2). These electron-dense sites of dermal-epidermal adhesion consisted of inner and outer hemidesmosome plaques associated with intracellular keratin intermediate filaments (Figure 3G-H, Figure S2). Beneath the plasma membrane of basal epidermal cells, the lamina lucida and lamina densa layers of the BM were each clearly distinguishable. Furthermore, the thickness of the lamina densa, where type IV collagen accumulates, was comparable between the skins of *mTurq2-Col4a1/+* and their WT littermates (Figure 3G-H). Thus, at the ultrastructural level, the skin of *mTurq2-Col4a1/+* embryos displayed all the hallmarks of normal dermal:epidermal junction assembly.

**Figure 3.**
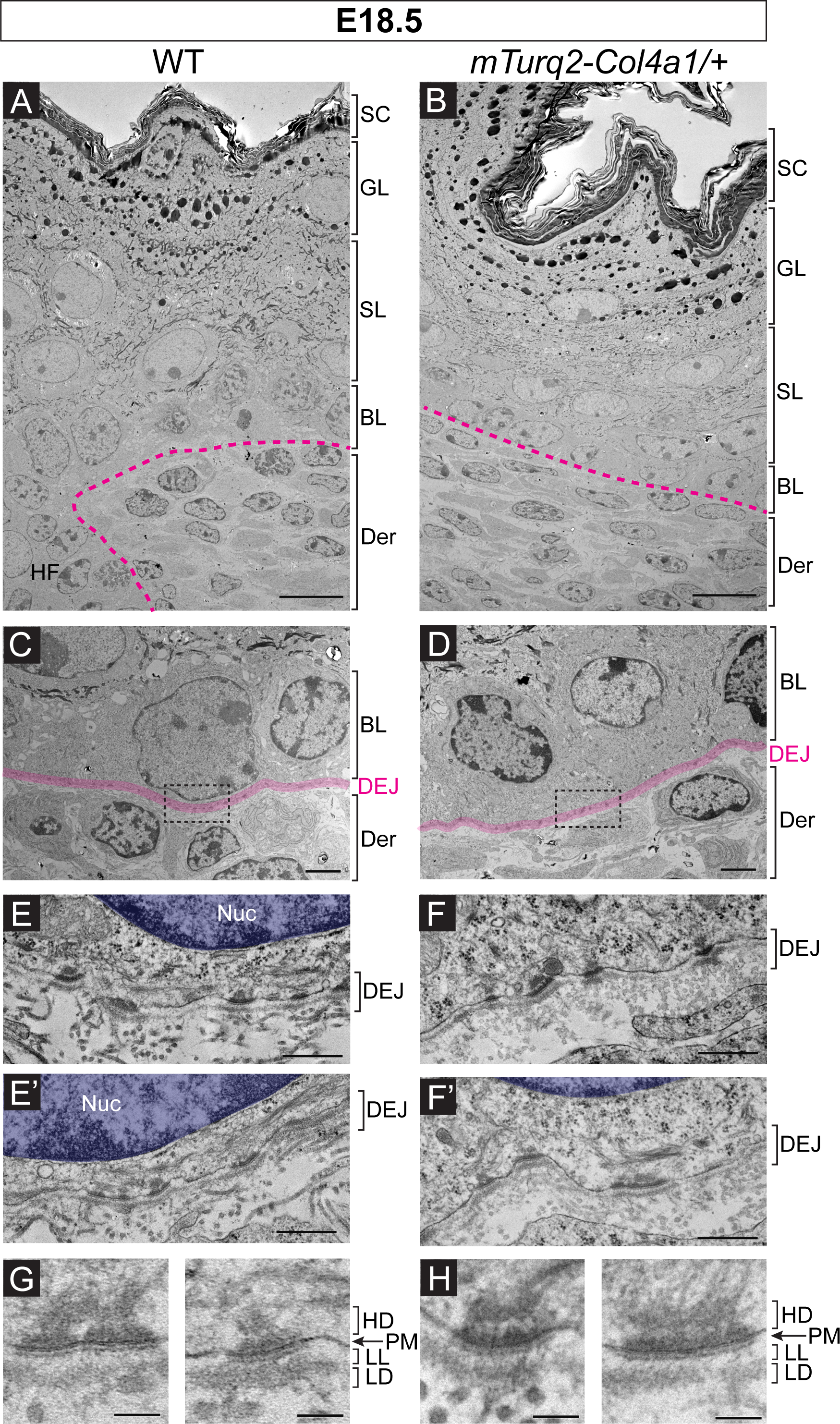
Normal ultrastructural organization of the epidermis and dermal-epidermal junction in *mTurq2-Col4a1/+* embryonic skin. Transmission electron micrographs of ultrathin skin sections from E18.5 WT control (A, C, E and G) and *mTurq2-Col4a1/+* (B, D, F, H) embryos. **(A-B)** Ultrastructural overview of skin architecture. Der = dermis, BL = basal layer, SL = spinous layer, GL = granular layer, SC = stratum corneum. Dotted line denotes dermal-epidermal boundary. Scale bars 10 µm. **(C-D)** Zoomed in view of the boundary between the basal epithelial layer and dermis. Dermal-epidermal junction (DEJ) is highlighted in pink. Dotted boxes denote regions magnified in (E) and (F). Scale bars 2 µm. **(E-F)** Two representative examples of DEJ region of the skin from WT (E and E’) and *mTurq2-Col4a1/+* (F and F’) embryos. Nuc=nucleus. Scale bars 500 nm. **(G-H)** Zoomed in views of individual hemidesmosomes at the DEJ. HD=hemidesmosome, PM = plasma membrane, BM = basement membrane. LL = lamina lucida, LD = lamina densa. Scale bars 100 nm.

### Characterization of mTurq2-COL4A1 expression in earlier embryonic stages

Having shown that our *mTurq2-Col4a1/+* mice display robust mTurq2-COL4A1 signal in both adult and late embryonic stage skin, we now sought to characterize the distribution of mTurq2-COL4A1 during BM assembly and maturation in earlier embryonic stages. To do this, we harvested embryos at E15.5 and labelled the backskins of both WT and *mTurq2-Col4a1/+* embryos with antibodies against laminin β-1 (Lamb1), collagen IV (ColIV) and perlecan (Figure 4). XZ optical reconstructions of Z stack images of E15.5 skins clearly show robust mTurq2-COL4A1 signal in the epidermal BM of both the IFE and developing hair follicles that recapitulates the ColIV antibody labelling (Figure 4B). As described above, we then calculated the PCC of the spatial relationship between Lamb1 and ColIV antibody labelling in both WT and *mTurq2-Col4a1/+* E15.5 epidermal BMs (Figure 4A-B) and detected no difference between the genotypes (Figure 4C), further validating our previous findings that incorporation of mTurq2-COL4A1 into the epidermal BM does not alter the interaction between these two supramolecular layers, at least in the skin.

**Figure 4.**
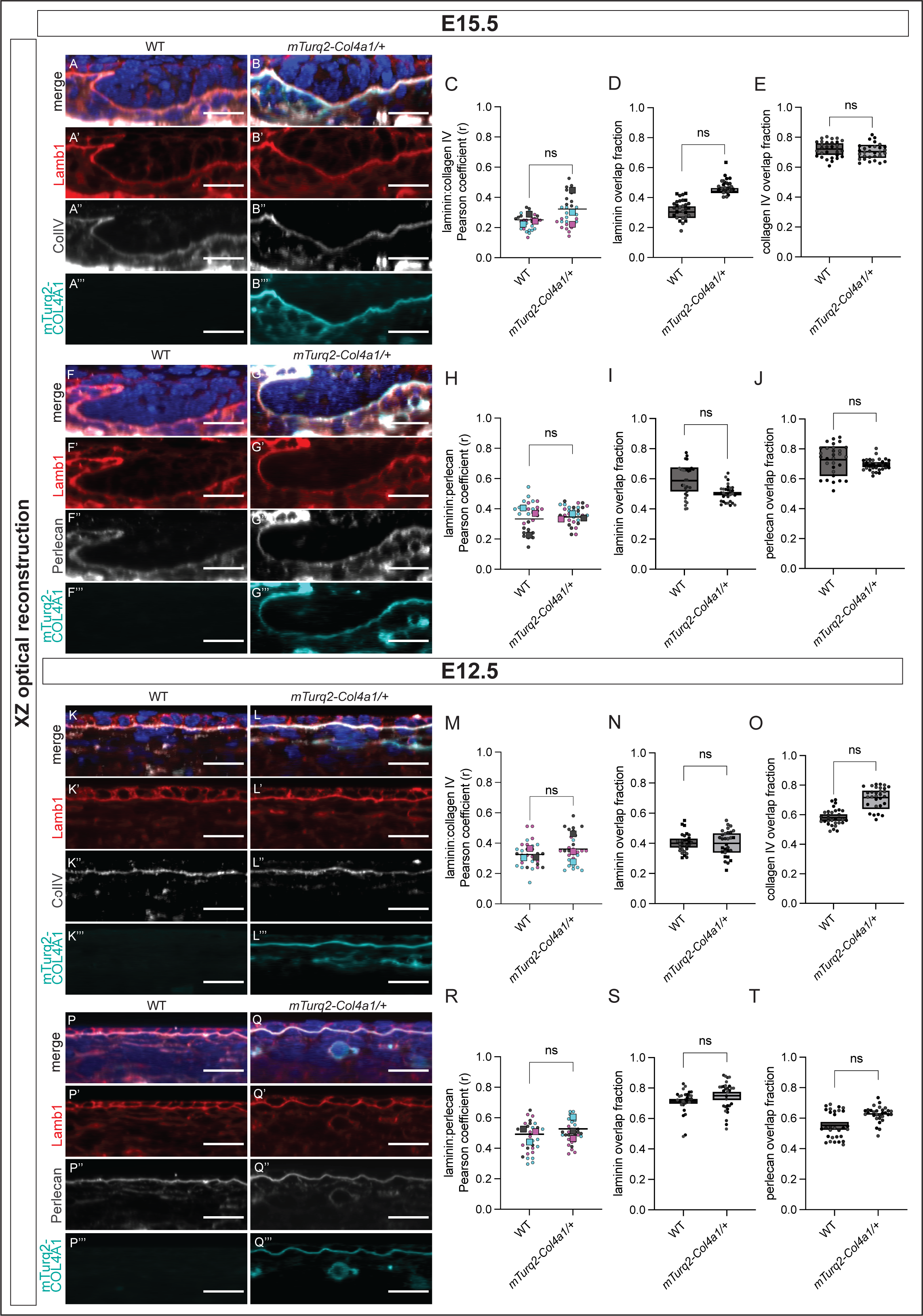
Spatial distribution and correlation between key basement membrane components in *mTurq2-Col4a1/+* embryonic backskins resembles that of WT littermates. **(A)** Average intensity projection (XZ optical reconstruction) of backskin from WT E15.5 embryo labelled with Hoechst (merge, top), anti-laminin β-1 antibody (A’) and anti-collagen IV antibody (A’’). No signal is observed in the mTurq2 channel (A’’’). **(B)** As (A) except backskin from an *mTurq2-Col4a1/+* embryo. Robust mTurq2-COL4A1 signal recapitulates the labelling pattern of the collagen IV antibody. **(C)** Pearson’s correlation coefficient (r) measuring the correlation between laminin β-1 and collagen IV expression (antibody labelling) in WT and *mTurq2-Col4a1/+* E15.5 backskins as labelled. Larger datapoints – biological replicate means (n=3 embryos), smaller datapoints – individual images per replicate (n=10 per embryo). **(D)** Fraction of laminin β-1 contribution to the laminin β-1:collagen IV overlap in E15.5 backskins; refer to main text for methodology. **(E)** As (D) except collagen IV contribution to laminin β-1:collagen IV overlap. **(F)** Average intensity projection (XZ optical reconstruction) of backskin from WT E15.5 embryo labelled with Hoechst (merge, top), anti-laminin β-1 antibody (F’) and anti-perlecan antibody (F’’). **(G)** As (F) except backskin from an *mTurq2-Col4a1/+* embryo. **(H-J)** As (C-E) except measuring the correlation and overlap fractions between laminin β-1 and perlecan. **(K-T)** As (A-J) except backskins are from E12.5 embryos. Scale bars 20 µm. ns = not significant, Mann-Whitney U-test.

To take this analysis further, we next measured how much of the laminin β-1 localization domain within the epidermal BM overlaps with that of the collagen IV localization domain. We analyzed high resolution images of the epidermal BM from both WT and *mTurq2-Col4a1/+* backskins and applied a custom FIJI macro to calculate the fraction of each protein that overlaps with the other (overlap fraction) (see Materials and Methods). Our results demonstrate that a higher fraction of collagen IV (∼70%) is found to overlap with laminin β-1 localization at E15.5 (Figure 4E). Laminin β-1 is more frequently observed not colocalizing with collagen IV (<50% overlap) (Figure 4D). Crucially, we observed no difference between the overlap fractions of laminin β-1 and collagen IV in WT and *mTurq2-Col4a1/+* (Figure 4D-E). We repeated these analyses with the laminin β-1 and perlecan localization domains (Figure 4H-J). Here we observed a similar contribution to the area of colocalization by both laminin β-1 and perlecan (Figure 4I-J), and again we saw no difference between WT and *mTurq2-Col4a1/+* in either analysis. As these results reflected what we observed in E18.5 embryos, we expanded our analysis to E12.5 embryos, a developmental time prior to both hair follicle development and stratification, when the basal progenitor layer is one-cell thick (Figure 4K-Q’’’). We hypothesized that due to significant growth of the embryo and expansion of the skin between these stages, the epidermal BM morphology and overlap composition would differ in E12.5 embryos. Again, we observed well-defined labelling of the epidermal BM with mTurq2-COL4A1 that recapitulated ColIV antibody labelling (Figure 4L”, L’’’) and detected no differences in the spatial relationships between laminin β-1 and collagen IV/perlecan in E12.5 WT and *mTurq2-Col4a1/+* BMs, respectively (Figure 4M-O and R-T). Surprisingly, we observed very similar overlap fractions of laminin β-1 and collagen IV/perlecan in E12.5 compared to E15.5. This suggests that the morphology and composition of the epidermal BM, in respect of these core components, is not changing significantly over time, and is established during early stages of embryonic skin development.

### Live imaging of *mTurq2-Col4a1/+* backskins by a Planar-Sagittal Multiview imaging approach reveals new insights into developmental BM dynamics

Having shown that the *mTurq2-Col4a1* mouse model faithfully labels the epidermal BM, we next explored its utility for live imaging developmental BM dynamics. As the epidermal BM has a thickness of around 100 nm and developing mouse skin is inherently undulating, to properly visualize BM dynamics a sagittal view of developing skin explants is necessary. Established live imaging approaches obtain planar XY images and the sagittal view is reconstructed from the Z-stack (Figure 5A). However, Abbe’s formula (lateral (XY) resolution = λ/2NA and axial (Z) resolution = 2λ/NA^2^) states that lateral resolution will always be greater than axial resolution in both widefield and confocal microscopy (Huszka and Gijs, 2019), so obtaining a high resolution sagittal image is a limitation of this method. To image the epidermal BM in all three dimensions with a similar resolution and without the need for software reconstruction, we developed a novel live imaging technique, Planar-Sagittal (PS) Multiview imaging. This simple technique involves folding the skin explant over an agarose ridge allowing the focal plane to be set at either the standard planar imaging plane (XY) or the sagittal imaging plane (‘XZ’) (Figure 5B and movie S1). This sagittal view of the developing skin allows for high resolution imaging of cells and tissues in ‘Z’, without the need for reconstruction. By combining PS Multiview imaging with *mTurq2-Col4a1/+* backskins in which all cell membranes are labelled with mTomato (*mTmG/mTmG*), we were able to long-term live image the epidermal BM in developing skin explants (Figure 5C and movie S1).

**Figure 5.**
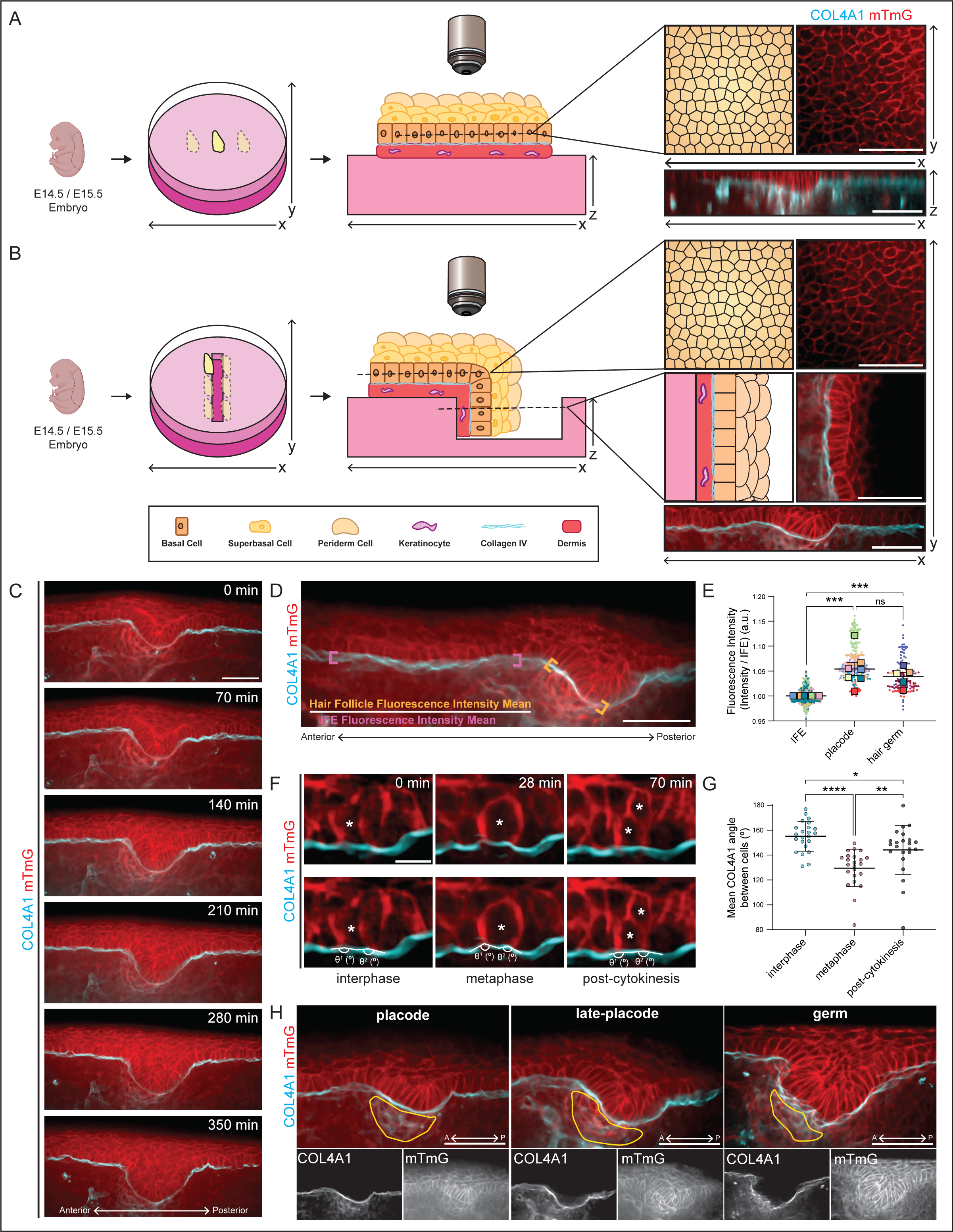
Planar-Sagittal (PS) Multiview imaging of *mTurq2-Col4a1* mouse model **(A)** Schematic of established live imaging method. The skin explant is mounted on an agarose pad and imaged using spinning disc microscopy. To visualize the tissue in Z, the image is software-reconstructed. Scale bar 50 µm. **(B)** Schematic of PS Multiview live imaging. The skin explant is mounted as previous, but over an agarose ridge. Both planar (XY) and sagittal (‘XZ’) focal planes are then imaged. Scale bar 50 µm. **(C)** Snapshots taken from a time-lapse live imaging movie of hair follicle invagination in E14.5 *mTurq2-Col4a1/+;mTmG/mTmG* backskin. **(D)** Representative image showing mean fluorescence intensity calculation methodology. Orange brackets highlight the hair follicle base region where signal intensity was measured. Magenta brackets highlight the IFE region where signal intensity was measured. Scale bar 50 µm. **(E)** Quantification of mTurq2-COL4A1 signal in IFE, placode, and hair germ stage. Each dot represents an individual fluorescence intensity measurement as shown in (D). Larger dots show means of independent experiments. Error bars = SEM. ***p=<0.001, Mann-Whitney U test. ns = not significant. **(F)** Snapshots taken from a time-lapse live imaging movie of basal cell division in E15.5 *mTurq2-Col4a1/+;mTmG/mTmG* backskin. Interphase, metaphase, and post-cytokinesis stages are shown (left to right). Bottom panel shows same images with membrane curvature measurements. Scale bar 10 µm. **(G)** Quantification of the average angle of BM deformation as shown in (F). Each dot represents an average of two angles. n=23 dividing cells. Error bars = mean+SD. *p=<0.05 (0.0366), **p=<0.01, ***p=<0.001, One-way ANOVA with Dunn’s multiple comparisons test. **(H)** Snapshots taken from time-lapse live imaging movies of placode, late-placode, and hair germ stages as labeled, in E14.5 and E15.5 *Col4a1/+;mTmG/mTmG* backskins. Bottom panel shows separate channels, note clear mTurq2-COL4A1 ‘baskets’ surrounding the developing dermal condensate. Scale bar 50 µm.

As we were able to now long-term live image the developing epidermal BM, we next wanted to visualize BM dynamics during the morphogenesis of hair follicles, which involves an initially thickened placode of basal epidermal cells that buds into the underlying dermis. The BM has been shown to play both an active and passive role during tissue development (Walma and Yamada, 2020), and is thought to be extensively remodeled during epithelial morphogenesis. For example, during branching morphogenesis, the BM surrounding actively growing branch tips becomes thin and/or discontinuous with microperforations, which is thought to allow for tip expansion and branch elongation (Harunaga et al., 2014, Spurlin et al., 2019, Sekiguchi and Yamada, 2018a). Given the wide-ranging roles of the BM during morphogenesis, we sought to investigate BM dynamics during hair follicle budding by live imaging E14.5-E15.5 *mTurq2-Col4a1/+;mTmG/mTmG* backskins over several hours using PS Multiview imaging (Figure 5C and movie S1).

We used the fluorescent intensity of the mTurq2-COL4A1 signal as a read-out for changes to the epidermal BM under an invaginating hair follicle relative to the neighboring IFE that is not undergoing significant morphological changes (Figure 5D). Surprisingly, the mTurq2-COL4A1 intensity was significantly greater at the actively growing base of developing hair follicles than at the IFE (Figure 5E). Although we had expected the intensity to become fainter, more diffuse, or discontinuous as downward budding occurred, the signal became brighter and qualitatively more uniform in appearance. This suggested that the epidermal BM may be compressed by the epithelium during hair follicle invagination, or that new COL4A1 incorporation exceeds proteolytic degradation during budding, unlike in other examples of budding/branching morphogenesis (Wang et al., 2017, Sekiguchi and Yamada, 2018a).

Another cellular force that acts upon the epidermal BM is cell division. Several studies have shown that when cells in culture approach mitosis, they detach from the ECM and disassemble their adhesions to retract from the matrix and round-up in metaphase (Champion et al., 2017, Suzuki and Takahashi, 2003, Yamakita et al., 1999). Similarly, epithelial cells are thought to reduce or lose their adhesions to the BM when undergoing division (Matus et al., 2014, Jones et al., 2019). Since basal cells within the epidermis divide frequently during development (Damen et al., 2021), we wanted to examine the interface between dividing basal cells and the epidermal BM. We analyzed several movies in which numerous basal cells were seen to go from interphase through to complete cytokinesis and to our surprise, we observed that when dividing cells rounded up in metaphase, they did not detach from the epidermal BM (Figure 5F, S4A-B, and movie S2). Instead, rounded cells remained closely adhered to the BM which subsequently deformed around the base of the dividing cell. To quantify this BM deformation, we measured the angle of the BM at the junction between the dividing cell and its neighbors before, during and after mitosis (interphase, metaphase, and cytokinesis, respectively). We found that the epidermal BM pinches inwards by an average of 25° underneath a rounded dividing cell and then reverts to the near-straight interphase angle after division, implicating maintained adhesion to the BM throughout mitosis (Figure 5G and S4C). In keeping with our previous observations of epidermal BM deformation underneath invaginating hair follicles, this suggests an inherent pliability of the epidermal BM during embryogenesis. BM stiffness has been shown to direct cellular behaviors and sculpt organs during development, and stiffer BMs are associated with cell migration and epithelial-to-mesenchymal transition (Crest et al., 2017, Ramos-Lewis and Page-McCaw, 2019, Wei et al., 2015). It is perhaps therefore intuitive that a BM underlying a rapidly proliferating progenitor cell layer during stratification needs to have sufficient pliability to allow cells to remain adhered throughout mitosis and to deform during morphogenesis.

Although we focused primarily on the epidermal BM for this study, PS Multiview imaging enabled visualization of a sharp delineation between the epidermis and dermis, with the dermis having a diffuse mTurq2-COL4A1 signal throughout which was completely absent in the epidermis, consistent with dermal fibroblasts being the major source of collagen IV in the skin (Zorina et al., 2023) (movie S1 and Figure 5H bottom panels). Moreover, several other structures in the dermis showed robust mTurq2-COL4A1 localization, including vascular BMs and as yet unidentified migrating cells containing bright puncta of mTurq2-COL4A1 (movie S1 and S3). We also noticed a consistent diffuse mTurq2-COL4A1 signal in the dermis that coincided with the dermal condensate, underlying each hair follicle. Upon closer inspection, it appeared that a ‘basket’ of collagen IV strands surrounds the cells of the dermal condensate as it develops (Figure 5H). This raises compelling questions regarding the role of collagen IV in maintaining the structural integrity of mesenchymal condensates during development, and it would be interesting to examine other mesenchymal condensates, such as those of the lung and kidney, to see if this phenomenon is conserved (Mammoto et al., 2015, Mammoto et al., 2011).

### Fluorescence Recovery After Photobleaching (FRAP) shows collagen IVa1 in the epidermal BM is highly stable

Having established that our *mTurq2-Col4a1* mouse model is amenable to live imaging approaches, we next assayed the stability of COL4A1 in the epidermal BM of *mTurq2-Col4a1/+* backskins at E13.5 and E15.5, stages when the skin undergoes significant growth and morphological changes (stratification and hair follicle development). Several studies in invertebrates have shown that COL4A1 within the BM is highly immobile (Morrissey et al., 2016, Matsubayashi et al., 2017, Keeley et al., 2020). The half-life of BM proteins in the *Drosophila* embryo has been measured to be on the order of ∼7-10 hours (Matsubayashi et al., 2020). Additionally, a recent study in mice found that active secretion of vascular BM COL4A1 could only be detected early in development, suggesting that secretion thereafter, throughout adult life, may occur at much lower levels (Lartey et al., 2023). We therefore hypothesized that COL4A1 would be highly stable in the epidermal BM but may show increased mobility during the earlier stage of E13.5 compared to E15.5, due to the rapid growth taking place during this time. We conducted FRAP on the inter-follicular epidermal BM at both E13.5 and E15.5 and observed minimal mobility, with only a small COL4A1 mobile fraction of <15 % at both stages (Figure 6C). We also performed FRAP on the BM around the rim of the hair follicle (E15.5) where the mTurq2-COL4A1 signal appears brightest (Figure 6A-B) and again observed minimal COL4A1 mobility. These data suggest that epidermal BM COL4A1 is highly stable within the epidermal BM, even as early as E13.5, consistent with other published datasets.

**Figure 6.**
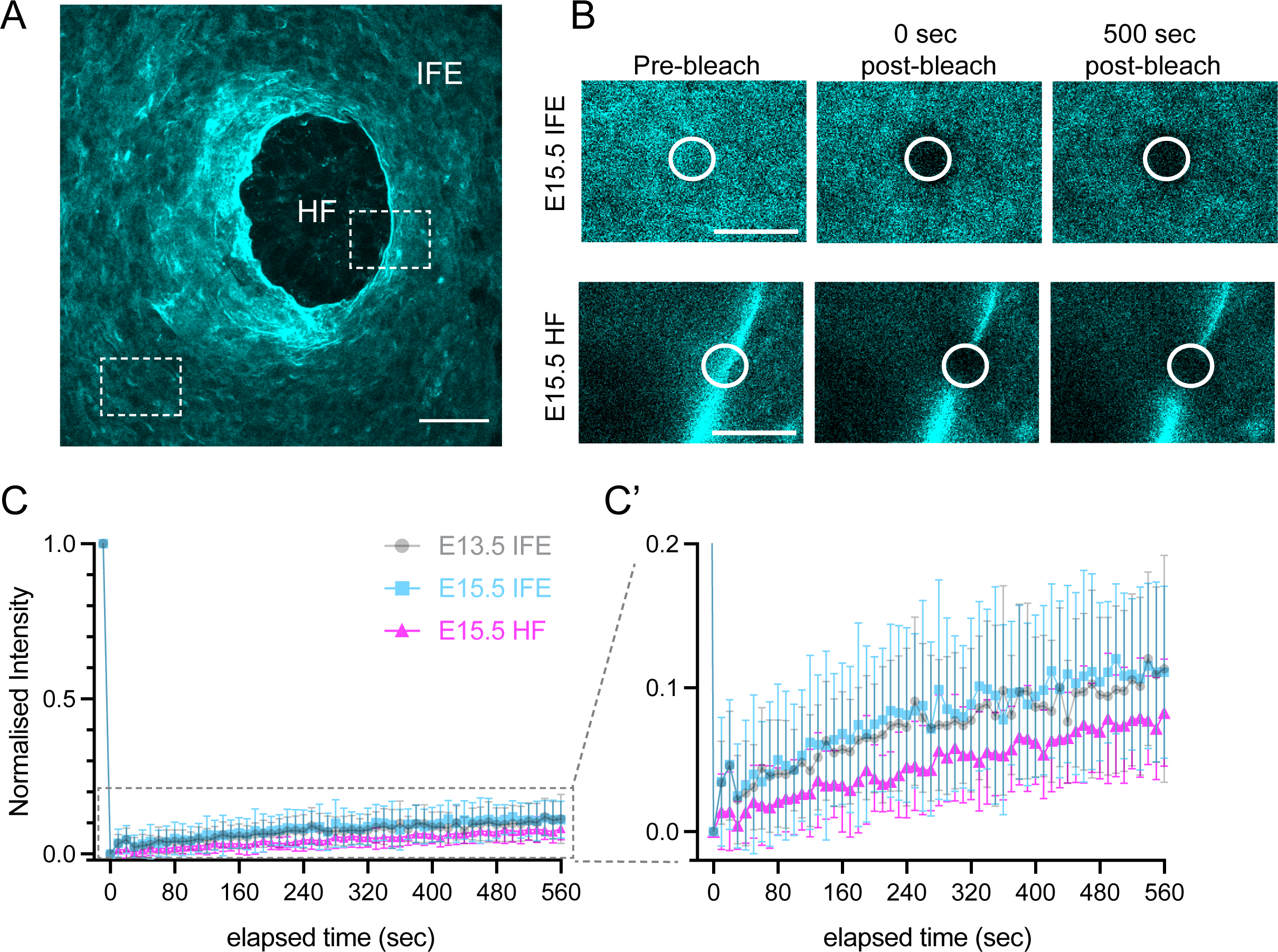
mTurq2-COL4A1 is highly stable within the embryonic epidermal basement membrane. **(A)** Planar overview of mTurq2-COL4A1 localization in E15.5 whole mount epidermis. White boxes indicate BM lying at the dermal-epidermal junction of the interfollicular epidermis (IFE), and along the rim of a budding hair follicle (HF). Scale bar 25 µm. See also Movie S3. **(B)** Still images from representative fluorescence recovery after photobleaching (FRAP) experiments of mTurq2-COL4A1 within the IFE (top panels) and rim of a hair follicle (bottom panels). Scale bars 10 µm. **(C)** FRAP recovery curves of mTurq2-COL4A1 within the IFE at E13.5 (grey circles; n=36), E15.5 (blue squares; n=32) and at the rim of E15.5 hair follicles (magenta triangles; n=30). Grey dotted box indicates region rescaled in (C’). Each point is the mean normalized intensity. Values were acquired across n>3 biological replicates. Error bars = SD.

## Discussion

Our understanding of the complexity of the BM has evolved significantly over the past few decades, and it is no longer considered a simple, static barrier between tissue compartments. Numerous studies have highlighted critical roles that the BM plays in many physiological processes, including development, cell signaling and disease progression (Töpfer, 2023, Pozzi et al., 2017, Randles et al., 2017, Banerjee et al., 2022). Elegant studies in *Drosophila* and *C.elegans* with fluorescently tagged BM components have significantly increased our understanding of the BM, in terms of its interaction with neighboring cells and the mobility of individual components within it (Isabella and Horne-Badovinac, 2015, Keeley et al., 2020, Kelley et al., 2019, Clay and Sherwood, 2015, Van De Bor et al., 2021). However, until recently, the key bottleneck in our understanding of mammalian, and indeed vertebrate BMs, has been the lack of models in which endogenous BM components are labelled. Several attempts to generate such models have been unsuccessful (Shaw et al., 2020), and *in vivo* studies have relied upon knock-in reporters which do not necessarily reflect endogenous protein expression and can be expressed ectopically and/or interfere with native protein function (Yamaguchi et al., 2022, Sztal et al., 2011, Futaki et al., 2023). The recent success by Morgner et al. (2023) in generating a *Lamb1-Dendra2* mouse model provided proof-of-principle that fluorescently tagging BM components in mouse is feasible, and in this study, we have advanced one step further by generating a mouse line in which the BM ubiquitous COL4A1 is labelled. Collagen IV is the primary component of all BMs, and we therefore envisage that our *mTurq2-Col4a1* mouse will open up new possibilities to study many different aspects of BM biology, from developmental morphogenesis and wound healing, through to cancer metastasis. Moreover, several human diseases are linked specifically to mutations in collagen IV genes. Mutations in *COL4A1* and *COL4A2* have been linked a wide range of human multi-system phenotypes, including ocular defects, porencephaly, cerebrovascular disease and myopathy, collectively termed Gould’s syndrome (Jeanne et al., 2012, Jeanne et al., 2015, Gould et al., 2005, Gould et al., 2006, Favor et al., 2007).

We focused on the epidermal BM for this study due to its accessibility and amenability to live imaging, and because little is known about epidermal BM dynamics during embryonic development. Using the novel *mTurq2-Col4a1* mouse model combined with an innovative PS Multiview imaging approach, we have been able to long-term live-image the developing epidermal BM in embryonic mouse skin for the first time. Our findings reveal changes in epidermal BM morphology during hair follicle morphogenesis and suggest that the BM may be compressed by the forces generated by hair follicle invagination. We show that significant BM deformation also occurs during cell division, suggesting an inherent pliability of the epidermal BM during embryogenesis. Moreover, our observations that dividing basal progenitor cells remain adhered to the BM during mitosis challenge the existing assumption that all dividing cells significantly reduce their adhesions to the BM, (Matus et al., 2014, Jones et al., 2019) and are in keeping with the integrin-rich contacts observed by Dix et al. (2018) during cell division. Our FRAP experiments provide proof-of-principle that the *mTurq2-Col4a1* mouse line can be used to study and quantify COL4A1 mobilty during assembly and our data agree with other studies demonstrating the highly stable nature of collagen IV within BMs, particularly the recent observations made by Larkey et al. (2023). The *Col4a1-P2A-eGFP* mouse line generated in their work labels cells that are actively secreting COL4A1 with eGFP. Because collagen IV is so stable, they wanted to determine when endothelial cells (EC) actively synthesize COL4A1 during development, as presence of COL4A1 in vascular BMs does not necessarily mean it is being actively secreted. They showed that secretion of COL4A1 by EC occurs in a temporally restricted manner during development, with eGFP expression (and thus COL4A1 secretion) being detected during the period E8.5-E16.5 but not by E18.5, or in any of the adult organs they observed (including skin). Although it is possible this observation reflects an inherent limitation of the mouse model, where subsequent COL4A1 secretion by EC is beyond the limits of detection, it is consistent with the idea that COL4A1, once established in a BM, is inherently very stable.

Importantly, although we focused on the epidermal BM in this study, the mTurq2-COL4A1 signal was observed in every physiological BM observed, in both embryos and adult tissues. Therefore, we foresee there is a great deal to learn about mammalian BM morphology, remodeling and mechanics that can be achieved using our *mTurq2-Col4a1* mouse.

A key limitation of this study is the homozygous lethality of the *mTurq2-Col4a1* mouse. Although we conducted several experiments to confirm the BMs in *mTurq2-Col4a1/+* animals are indistinguishable from WT littermates, we have not been able to explain the lethality of the homozygous genotype. For studies of collagen IV protein biochemistry therefore, the *mTurq2-Col4a1* mouse may not be the best suited model. One possible explanation for the absence of homozygous animals is that heterotrimers containing two mTurq2-COL4A1 molecules (collagen α1/α1/α2 (IV)) may cause problems with secretion; mutant COL4A1 or COL4A2 has been shown to accumulate within cells, reducing secretion and leading to extracellular deficiency (Jeanne et al., 2015). Inhibited secretion was also observed with C-terminal tagged human laminin β-1 in adenocarcinoma cells (Shaw et al., 2020). Alternatively, given that the 7S domain is important for crosslinking collagen IV into a lattice network, it may be that two mTurq2 molecules per collagen IV trimer interferes with 7S-mediated crosslinking, but one mTurq2 per trimer can be tolerated. Nevertheless, as the mTurq2-COL4A1 signal is bright and photostable in heterozygous *mTurq2-Col4a1/+* mice, this limitation does not impede the use of our mouse line to visualize BM dynamics live, *in vivo*. Furthermore, the *Lamb1-Dendra2* mouse, currently the only other mouse line in which a BM component is tagged at the endogenous level, is also largely homozygous lethal (Morgner et al., 2023). Finally, mTurquoise2 has an excitation/emission spectrum of 434/474 nm (Goedhart et al., 2012), necessitating the use of a 445 nm laser for optimal imaging results, which is often not included as part of the traditional 4-color laser confocal set-up.

Using a novel *mTurq2-Col4a1* mouse model combined with an innovative PS Multiview imaging technique, we have visualized previously unobserved dynamics of developing skin. Our findings reveal unique insights into the epidermal BM during mouse embryonic development and highlight its inherent pliability in the context of morphological change. Although the large degree of growth, morphogenetic movement and cell division occurring during the period studied in this work suggested a need for the collagen IV supramolecular network to be extensively degraded, our observations show that once the collagen IV network is established early in development it appears strikingly stable, which may be due to either an equilibrium of degradation and secretion or an inherent pliability requiring little degradation to accommodate morphogenetic movements. Even during periods of rapid epithelial proliferation and morphogenetic change, mTurq2-COL4A1 within the epidermal BM is remarkably stable, showing minimal mobility. These data suggest that the observed pliability of the structure is a physical attribute of a relatively stable, but flexible meshwork. Given the close association between increases in ECM stiffness, aging, cancer metastasis and disease progression, our data raise questions about the behavioral changes cells might exhibit on a more rigid BM that can no longer yield to cellular forces. With the *mTurq2-Col4a1* mouse model in hand, these questions are now feasible to address.

## Acknowledgements

The authors thank the staff of the Princeton Laboratory Animal Resources (LAR) and the Princeton Core Imaging Facility (CIF), particularly Dr. Gary S. Laevsky and Dr. Sha Wang. Many useful discussions were had with Professor Doug Gould (UCSF), Dr. Jessica Morgner (Netherlands Cancer Institute) and Professor Dorus Gadella (University of Amsterdam), and the authors extend our heartfelt thanks. This work was supported by NIH/NICHD grant R01HD105009 (D.D.), NIH/NIAMS grant R01AR068320 (D.D.), NIH NIGMS T32GM007388 (B.T.), the New Jersey Alliance for Clinical and Translational Science (D.D., R.J, B.T.) and the Ludwig Institute for Cancer Research (D.D. and R.J.).

## Author contributions

R.J. and D.D. conceptualized the study; R.J. designed and performed most experiments. B.T. designed the PS Multiview imaging methodology and performed live-imaging experiments with R.J. Samples prepared for EM by R.J. were further prepared and EM imaged by H.P. The mouse model was generated with the help of B.J. and E.P. P.S. performed FRAP experiments and analysis with R.J.; K.L. also performed experiments. R.J. and D.D. wrote the manuscript with all authors having read it and provided comments.

## Notes

### Competing Interest Statement

The authors have declared no competing interest.

## References

Albini, A. & Noonan, D. M. 2010. The ‘chemoinvasion’ assay, 25 years and still going strong: the use of reconstituted basement membranes to study cell invasion and angiogenesis. Current Opinion in Cell Biology, 22, 677–689.

Bahr, J. C., Li, X.-Y., Feinberg, T. Y., Jiang, L. & Weiss, S. J. 2022. Divergent regulation of basement membrane trafficking by human macrophages and cancer cells. Nature Communications, 13, 6409.

Banerjee, S., ISAACMAN-Beck, J., Schneider, V. A. & Granato, M. 2013. A novel role for Lh3 dependent ECM modifications during neural crest cell migration in zebrafish. PloS one, 8, e54609.

Banerjee, S., Lo, W.-C., Majumder, P., Roy, D., Ghorai, M., Shaikh, N. K., Kant, N., Shekhawat, M. S., Gadekar, V. S. & Ghosh, S. 2022. Multiple roles for basement membrane proteins in cancer progression and EMT. European Journal of Cell Biology, 101, 151220.

Behrens, D. T., Villone, D., Koch, M., Brunner, G., Sorokin, L., Robenek, H., BRUCKNER-Tuderman, L., Bruckner, P. & Hansen, U. 2012. The epidermal basement membrane is a composite of separate laminin- or collagen IV-containing networks connected by aggregated perlecan, but not by nidogens. J Biol Chem, 287, 18700–9.

Buszczak, M., Paterno, S., Lighthouse, D., Bachman, J., Planck, J., Owen, S., Skora, A. D., Nystul, T. G., Ohlstein, B., Allen, A., Wilhelm, J. E., Murphy, T. D., Levis, R. W., Matunis, E., Srivali, N., Hoskins, R. A. & Spradling, A. C. 2007. The carnegie protein trap library: a versatile tool for Drosophila developmental studies. Genetics, 175, 1505–31.

Cetera, M., Leybova, L., Joyce, B. & Devenport, D. 2018. Counter-rotational cell flows drive morphological and cell fate asymmetries in mammalian hair follicles. Nat Cell Biol, 20, 541–552.

Champion, L., Linder, M. I. & Kutay, U. 2017. Cellular reorganization during mitotic entry. Trends in cell biology, 27, 26–41.

Chan, A. & Mauro, T. 2011. Acidification in the epidermis and the role of secretory phospholipases. Dermato-endocrinology, 3, 84–90.

Chlasta, J., Milani, P., Runel, G., Duteyrat, J.-L., Arias, L., Lamiré, L.-A., Boudaoud, A. & Grammont, M. 2017. Variations in basement membrane mechanics are linked to epithelial morphogenesis. Development, 144, 4350–4362.

Clay, M. R. & Sherwood, D. R. 2015. Basement membranes in the worm: a dynamic scaffolding that instructs cellular behaviors and shapes tissues. Current topics in membranes, 76, 337–371.

Concordet, J.-P. & Haeussler, M. 2018. CRISPOR: intuitive guide selection for CRISPR/Cas9 genome editing experiments and screens. Nucleic Acids Research, 46, W242–W245.

Costa, D. S., KENNY-Ganzert, I. W., Chi, Q., Park, K., Kelley, L. C., Garde, A., Matus, D. Q., Park, J., Yogev, S. & Goldstein, B. 2023. The Caenorhabditis elegans anchor cell transcriptome: ribosome biogenesis drives cell invasion through basement membrane. Development, 150, dev201570.

Costantini, L. M., Fossati, M., Francolini, M. & Snapp, E. L. 2012. Assessing the Tendency of Fluorescent Proteins to Oligomerize Under Physiologic Conditions. Traffic, 13, 643–649.

Crest, J., DIZ-Munoz, A., Chen, D.-Y., Fletcher, D. A. & Bilder, D. 2017. Organ sculpting by patterned extracellular matrix stiffness. Elife, 6, e24958.

Damen, M., Wirtz, L., Soroka, E., Khatif, H., Kukat, C., Simons, B. D. & Bazzi, H. 2021. High proliferation and delamination during skin epidermal stratification. Nature Communications, 12, 3227.

Denda, M., Hosoi, J. & Asida, Y. 2000. Visual imaging of ion distribution in human epidermis. Biochem Biophys Res Commun, 272, 134–7.

Dix, C. L., Matthews, H. K., Uroz, M., Mclaren, S., Wolf, L., Heatley, N., Win, Z., Almada, P., Henriques, R. & Boutros, M. 2018. The role of mitotic cell-substrate adhesion remodeling in animal cell division. Developmental cell, 45, 132–145. e3.

Favor, J., Gloeckner, C. J., Janik, D., Klempt, M., NEUHÄUSER-Klaus, A., Pretsch, W., Schmahl, W. & QUINTANILLA-Fend, L. 2007. Type IV procollagen missense mutations associated with defects of the eye, vascular stability, the brain, kidney function and embryonic or postnatal viability in the mouse, Mus musculus: an extension of the Col4a1 allelic series and the identification of the first two Col4a2 mutant alleles. Genetics, 175, 725–736.

Frantz, C., Stewart, K. M. & Weaver, V. M. 2010. The extracellular matrix at a glance. J Cell Sci, 123, 4195–200.

Futaki, S., Horimoto, A., Shimono, C., Norioka, N., Taniguchi, Y., Hamaoka, H., Kaneko, M., Shigeta, M., Abe, T. & Sekiguchi, K. 2023. Visualization of basement membranes by a nidogen-based fluorescent reporter in mice. Matrix Biology Plus, 18, 100133.

Glentis, A., Oertle, P., Mariani, P., Chikina, A., EL Marjou, F., Attieh, Y., Zaccarini, F., Lae, M., Loew, D. & Dingli, F. 2017. Cancer-associated fibroblasts induce metalloprotease-independent cancer cell invasion of the basement membrane. Nature communications, 8, 924.

Goedhart, J., VON Stetten, D., NOIRCLERC-Savoye, M., Lelimousin, M., Joosen, L., Hink, M. A., VAN Weeren, L., Gadella, T. W. J. & Royant, A. 2012. Structure-guided evolution of cyan fluorescent proteins towards a quantum yield of 93%. Nature Communications, 3, 751.

Gould, D. B., Phalan, F. C., Breedveld, G. J., VAN Mil, S. E., Smith, R. S., Schimenti, J. C., Aguglia, U., VAN DER Knaap, M. S., Heutink, P. & John, S. W. 2005. Mutations in Col4a1 cause perinatal cerebral hemorrhage and porencephaly. Science, 308, 1167–1171.

Gould, D. B., Phalan, F. C., VAN Mil, S. E., Sundberg, J. P., Vahedi, K., Massin, P., Bousser, M. G., Heutink, P., Miner, J. H. & Tournier-Lasserve, E. 2006. Role of COL4A1 in small-vessel disease and hemorrhagic stroke. New England Journal of Medicine, 354, 1489–1496.

Gu, B., Posfai, E., Gertsenstein, M. & Rossant, J. 2020. Efficient Generation of Large-Fragment Knock-In Mouse Models Using 2-Cell (2C)-Homologous Recombination (HR)-CRISPR. Current Protocols in Mouse Biology, 10, e67.

Gu, B., Posfai, E. & Rossant, J. 2018. Efficient generation of targeted large insertions by microinjection into two-cell-stage mouse embryos. Nat Biotechnol, 36, 632–637.

Gunwar, S., Ballester, F., Noelken, M. E., Sado, Y., Ninomiya, Y. & Hudson, B. G. 1998. Glomerular basement membrane: identification of a novel disulfide-cross-linked network of α3, α4, and α5 chains of type IV collagen and its implications for the pathogenesis of Alport syndrome. Journal of Biological Chemistry, 273, 8767–8775.

Guo, X., Johnson, J. J. & Kramer, J. M. 1991. Embryonic lethality caused by mutations in basement membrane collagen of C. elegans. Nature, 349, 707–709.

Guo, X. & Kramer, J. 1989. The two Caenorhabditis elegans basement membrane (type IV) collagen genes are located on separate chromosomes. Journal of Biological Chemistry, 264, 17574–17582.

Harmansa, S., Erlich, A., Eloy, C., Zurlo, G. & Lecuit, T. 2023. Growth anisotropy of the extracellular matrix shapes a developing organ. Nature Communications, 14, 1220.

Harunaga, J. S., Doyle, A. D. & Yamada, K. M. 2014. Local and global dynamics of the basement membrane during branching morphogenesis require protease activity and actomyosin contractility. Developmental biology, 394, 197–205.

Hohenester, E. & Yurchenco, P. D. 2013. Laminins in basement membrane assembly. Cell adhesion & migration, 7, 56–63.

Huszka, G. & Gijs, M. A. 2019. Super-resolution optical imaging: A comparison. Micro and Nano Engineering, 2, 7–28.

Hynes, R. O. 1992. Integrins: versatility, modulation, and signaling in cell adhesion. Cell, 69, 11–25.

Isabella, A. J. & HORNE-Badovinac, S. 2015. Dynamic regulation of basement membrane protein levels promotes egg chamber elongation in Drosophila. Developmental biology, 406, 212–221.

Jeanne, M., Jorgensen, J. & Gould, D. B. 2015. Molecular and genetic analyses of collagen type IV mutant mouse models of spontaneous intracerebral hemorrhage identify mechanisms for stroke prevention. Circulation, 131, 1555–1565.

Jeanne, M., LABELLE-Dumais, C., Jorgensen, J., Kauffman, W. B., Mancini, G. M., Favor, J., Valant, V., Greenberg, S. M., Rosand, J. & Gould, D. B. 2012. COL4A2 mutations impair COL4A1 and COL4A2 secretion and cause hemorrhagic stroke. The American Journal of Human Genetics, 90, 91–101.

Jones, F. E., Bailey, M. A., Murray, L. S., Lu, Y., Mcneilly, S., SCHLÖTZER-Schrehardt, U., Lennon, R., Sado, Y., Brownstein, D. G. & Mullins, J. J. 2016. ER stress and basement membrane defects combine to cause glomerular and tubular renal disease resulting from Col4a1 mutations in mice. Disease models & mechanisms, 9, 165–176.

Jones, M. C., Zha, J. & Humphries, M. J. 2019. Connections between the cell cycle, cell adhesion and the cytoskeleton. Philosophical Transactions of the Royal Society B, 374, 20180227.

Keeley, D. P., Hastie, E., Jayadev, R., Kelley, L. C., Chi, Q., Payne, S. G., Jeger, J. L., Hoffman, B. D. & Sherwood, D. R. 2020. Comprehensive Endogenous Tagging of Basement Membrane Components Reveals Dynamic Movement within the Matrix Scaffolding. Dev Cell, 54, 60–74 e7.

Kelley, L. C., Chi, Q., Caceres, R., Hastie, E., Schindler, A. J., Jiang, Y., Matus, D. Q., Plastino, J. & Sherwood, D. R. 2019. Adaptive F-Actin Polymerization and Localized ATP Production Drive Basement Membrane Invasion in the Absence of MMPs. Dev Cell, 48, 313–328 e8.

Kelley, L. C., Lohmer, L. L., Hagedorn, E. J. & Sherwood, D. R. 2014. Traversing the basement membrane in vivo: a diversity of strategies. J Cell Biol, 204, 291–302.

Kelley, L. C., Wang, Z., Hagedorn, E. J., Wang, L., Shen, W., Lei, S., Johnson, S. A. & Sherwood, D. R. 2017. Live-cell confocal microscopy and quantitative 4D image analysis of anchor-cell invasion through the basement membrane in Caenorhabditis elegans. nature protocols, 12, 2081–2096.

KENNY-Ganzert, I. W. & Sherwood, D. R. 2023. The C. elegans anchor cell: A model to elucidate mechanisms underlying invasion through basement membrane. Seminars in Cell & Developmental Biology.

Kruegel, J. & Miosge, N. 2010. Basement membrane components are key players in specialized extracellular matrices. Cellular and Molecular Life Sciences, 67, 2879–2895.

Lambert, T. J. 2019. FPbase: a community-editable fluorescent protein database. Nature Methods, 16, 277–278.

Lartey, N. L., VAN DER Ent, M., Alonzo, R., Chen, D. & King, P. D. 2023. A temporally-restricted pattern of endothelial cell collagen 4 alpha 1 expression during embryonic development determined with a novel knockin Col4a1-P2A-eGFP mouse line. *Genesis*, e23539.

Lebleu, V. S., Macdonald, B. & Kalluri, R. 2007. Structure and Function of Basement Membranes. Experimental Biology and Medicine, 232, 1121–1129.

Leonard, C. E. & Taneyhill, L. A. 2020. The road best traveled: Neural crest migration upon the extracellular matrix. Seminars in Cell & Developmental Biology, 100, 177–185.

M, M. P. 1992. Basement Membrane Proteins: Structure, Assembly, and Cellular Interactions. Critical Reviews in Biochemistry and Molecular Biology, 27, 93–127.

Ma, M., Cao, X., Dai, J. & PASTOR-Pareja, J. C. 2017. Basement membrane manipulation in Drosophila wing discs affects Dpp retention but not growth mechanoregulation. Developmental cell, 42, 97–106. e4.

Mak, K. M. & Mei, R. 2017. Basement Membrane Type IV Collagen and Laminin: An Overview of Their Biology and Value as Fibrosis Biomarkers of Liver Disease. The Anatomical Record, 300, 1371–1390.

Mammoto, T., Mammoto, A., Jiang, A., Jiang, E., Hashmi, B. & Ingber, D. E. 2015. Mesenchymal condensation-dependent accumulation of collagen VI stabilizes organ-specific cell fates during embryonic tooth formation. Developmental Dynamics, 244, 713–723.

Mammoto, T., Mammoto, A., Torisawa, Y.-S., Tat, T., Gibbs, A., Derda, R., Mannix, R., De Bruijn, M., Yung, C. W. & Huh, D. 2011. Mechanochemical control of mesenchymal condensation and embryonic tooth organ formation. Developmental cell, 21, 758–769.

Matsubayashi, Y., Louani, A., Dragu, A., SANCHEZ-Sanchez, B. J., SERNA-Morales, E., Yolland, L., Gyoergy, A., Vizcay, G., Fleck, R. A., Heddleston, J. M., Chew, T. L., Siekhaus, D. E. & Stramer, B. M. 2017. A Moving Source of Matrix Components Is Essential for De Novo Basement Membrane Formation. Curr Biol, 27, 3526–3534 e4.

Matsubayashi, Y., SANCHEZ-Sanchez, B. J., Marcotti, S., SERNA-Morales, E., Dragu, A., VIZCAY-Barrena, G., Fleck, R. A. & Stramer, B. M. 2020. Rapid homeostatic turnover of embryonic ECM during tissue morphogenesis. Developmental Cell, 54, 33–42. e9.

Matus, D. Q., Chang, E., MAKOHON-Moore, S. C., Hagedorn, M. A., Chi, Q. & Sherwood, D. R. 2014. Cell division and targeted cell cycle arrest opens and stabilizes basement membrane gaps. Nature communications, 5, 4184.

Micalizzi, D. S., Farabaugh, S. M. & Ford, H. L. 2010. Epithelial-mesenchymal transition in cancer: parallels between normal development and tumor progression. Journal of mammary gland biology and neoplasia, 15, 117–134.

Morgner, J., Bornes, L., Hahn, K., LOPEZ-Iglesias, C., Kroese, L., Pritchard, C. E. J., Vennin, C., Peters, P. J., Huijbers, I. & VAN Rheenen, J. 2023. A Lamb1Dendra2 mouse model identifies basement-membrane-producing origins and dynamics in PyMT breast tumors. Dev Cell, 58, 535–549 e5.

Morin, X., Daneman, R., Zavortink, M. & Chia, W. 2001. A protein trap strategy to detect GFP-tagged proteins expressed from their endogenous loci in Drosophila. Proc Natl Acad Sci U S A, 98, 15050–5.

Morrissey, M. A., Jayadev, R., Miley, G. R., Blebea, C. A., Chi, Q., Ihara, S. & Sherwood, D. R. 2016. SPARC Promotes Cell Invasion In Vivo by Decreasing Type IV Collagen Levels in the Basement Membrane. PLoS Genet, 12, e1005905.

Morrissey, M. A. & Sherwood, D. R. 2015. An active role for basement membrane assembly and modification in tissue sculpting. Journal of cell science, 128, 1661–1668.

Murray, L. S., Lu, Y., Taggart, A., VAN Regemorter, N., Vilain, C., Abramowicz, M., Kadler, K. E. & VAN Agtmael, T. 2014. Chemical chaperone treatment reduces intracellular accumulation of mutant collagen IV and ameliorates the cellular phenotype of a COL4A2 mutation that causes haemorrhagic stroke. Human molecular genetics, 23, 283–292.

Nakaya, Y. & Sheng, G. 2013. EMT in developmental morphogenesis. Cancer letters, 341, 9–15.

Netzer, K.-O., Suzuki, K., Itoh, Y., Hudson, B. G. & Khalifah, R. G. 1998. Comparative analysis of the noncollagenous NC1 domain of type IV collagen: Identification of structural features important for assembly, function, and pathogenesis. Protein Science, 7, 1340–1351.

Özbek, S., Balasubramanian, P. G., CHIQUET-Ehrismann, R., Tucker, R. P. & Adams, J. C. 2010. The evolution of extracellular matrix. Molecular biology of the cell, 21, 4300–4305.

PAGE-Mccaw, A. Remodeling the model organism: matrix metalloproteinase functions in invertebrates. Seminars in cell & developmental biology, 2008. Elsevier, 14–23.

PAGE-Mccaw, A., Ewald, A. J. & Werb, Z. 2007. Matrix metalloproteinases and the regulation of tissue remodelling. Nature reviews Molecular cell biology, 8, 221–233.

Pöschl, E., SCHLÖTZER-Schrehardt, U., Brachvogel, B., Saito, K., Ninomiya, Y. & Mayer, U. 2004. Collagen IV is essential for basement membrane stability but dispensable for initiation of its assembly during early development. Development, 131, 1619–1628.

Pozzi, A., Yurchenco, P. D. & Iozzo, R. V. 2017. The nature and biology of basement membranes. Matrix Biology, 57-58, 1–11.

RAMOS-Lewis, W. & PAGE-Mccaw, A. 2019. Basement membrane mechanics shape development: Lessons from the fly. Matrix Biology, 75, 72–81.

Randles, M. J., Humphries, M. J. & Lennon, R. 2017. Proteomic definitions of basement membrane composition in health and disease. Matrix Biology, 57-58, 12–28.

Ratzinger, G., Stoitzner, P., Ebner, S., Lutz, M. B., Layton, G. T., Rainer, C., Senior, R. M., Shipley, J. M., Fritsch, P. & Schuler, G. 2002. Matrix metalloproteinases 9 and 2 are necessary for the migration of Langerhans cells and dermal dendritic cells from human and murine skin. The Journal of Immunology, 168, 4361–4371.

Rei, K., Hiroaki, T., Sheng, D., Yasuyuki, F. & Yoichiro, T. 2021. Tumor-Cell Invasion Initiates at Invasion Hotspots, an Epithelial Tissue-Intrinsic Microenvironment. bioRxiv, 2021.09.28.462102.

Schindler, A. J. & Sherwood, D. R. 2013. Morphogenesis of the Caenorhabditis elegans vulva. Wiley Interdisciplinary Reviews: Developmental Biology, 2, 75–95.

Schwarzbauer, J. 1999. Basement membrane: Putting up the barriers. Current Biology, 9, R242–R244.

Sekiguchi, R. & Yamada, K. M. 2018a. Basement membranes in development and disease. Current topics in developmental biology, 130, 143–191.

Sekiguchi, R. & Yamada, K. M. 2018b. Chapter Four - Basement Membranes in Development and Disease. In: Litscher, E. S. & Wassarman, P. M. (eds.) Current Topics in Developmental Biology. Academic Press.

Shaw, L., Williams, R. & Hamill, K. 2020. CRISPR-Cas9-mediated labelling of the C-terminus of human laminin β1 leads to secretion inhibition. BMC Research Notes, 13, 1–7.

Sherwood, D. R. 2021. Basement membrane remodeling guides cell migration and cell morphogenesis during development. Curr Opin Cell Biol, 72, 19–27.

Sherwood, D. R. & Sternberg, P. W. 2003. Anchor cell invasion into the vulval epithelium in C. elegans. Developmental cell, 5, 21–31.

Spurlin, J. W., Siedlik, M. J., Nerger, B. A., Pang, M.-F., Jayaraman, S., Zhang, R. & Nelson, C. M. 2019. Mesenchymal proteases and tissue fluidity remodel the extracellular matrix during airway epithelial branching in the embryonic avian lung. Development, 146, dev175257.

Stanley, J. R., Woodley, D. T., Katz, S. I. & Martin, G. R. 1982. Structure and Function of Basement Membrane. Journal of Investigative Dermatology, 79, 69–72.

Suzuki, K. & Takahashi, K. 2003. Reduced cell adhesion during mitosis by threonine phosphorylation of β1 integrin. Journal of cellular physiology, 197, 297–305.

Sztal, T., Berger, S., Currie, P. D. & Hall, T. E. 2011. Characterization of the laminin gene family and evolution in zebrafish. Developmental dynamics, 240, 422–431.

Tamori, Y., Suzuki, E. & Deng, W.-M. 2016. Epithelial Tumors Originate in Tumor Hotspots, a Tissue-Intrinsic Microenvironment. PLOS Biology, 14, e1002537.

Timpl, R. 1989. Structure and biological activity of basement membrane proteins. European Journal of Biochemistry, 180, 487–502.

Timpl, R. 1996. Macromolecular organization of basement membranes. Current Opinion in Cell Biology, 8, 618–624.

Töpfer, U. 2023. Basement membrane dynamics and mechanics in tissue morphogenesis. Biology Open, 12.

VAN DE Bor, V., Loreau, V., Malbouyres, M., Cerezo, D., Placenti, A., Ruggiero, F. & Noselli, S. 2021. A dynamic and mosaic basement membrane controls cell intercalation in Drosophila ovaries. Development, 148, dev195511.

Walma, D. A. C. & Yamada, K. M. 2020. The extracellular matrix in development. Development, 147, dev175596.

Wang, S., Sekiguchi, R., Daley, W. P. & Yamada, K. M. 2017. Patterned cell and matrix dynamics in branching morphogenesis. Journal of Cell Biology, 216, 559–570.

Wei, S. C., Fattet, L., Tsai, J. H., Guo, Y., Pai, V. H., Majeski, H. E., Chen, A. C., Sah, R. L., Taylor, S. S. & Engler, A. J. 2015. Matrix stiffness drives epithelial–mesenchymal transition and tumour metastasis through a TWIST1–G3BP2 mechanotransduction pathway. Nature cell biology, 17, 678–688.

Yamaguchi, N., Zhang, Z., Schneider, T., Wang, B., Panozzo, D. & Knaut, H. 2022. Rear traction forces drive adherent tissue migration in vivo. Nature Cell Biology, 24, 194–204.

Yamakita, Y., Totsukawa, G., Yamashiro, S., Fry, D., Zhang, X., Hanks, S. K. & Matsumura, F. 1999. Dissociation of FAK/p130CAS/c-Src complex during mitosis: role of mitosis-specific serine phosphorylation of FAK. The Journal of cell biology, 144, 315–324.

Yasunaga, K.-I., Kanamori, T., Morikawa, R., Suzuki, E. & Emoto, K. 2010. Dendrite reshaping of adult Drosophila sensory neurons requires matrix metalloproteinase-mediated modification of the basement membranes. Developmental cell, 18, 621–632.

Yurchenco, P. D. 2011. Basement membranes: cell scaffoldings and signaling platforms. Cold Spring Harb Perspect Biol, 3.

Yurchenco, P. D., Amenta, P. S. & Patton, B. L. 2004. Basement membrane assembly, stability and activities observed through a developmental lens. Matrix Biology, 22, 521–538.

Zorina, A., Zorin, V., Isaev, A., Kudlay, D., Vasileva, M. & Kopnin, P. 2023. Dermal Fibroblasts as the Main Target for Skin Anti-Age Correction Using a Combination of Regenerative Medicine Methods. Current Issues in Molecular Biology [Online], 45.

