## Supplementary material for "A Window into Mammalian Basement Membrane Development: Insights from the *mTurq2-Col4a1* Mouse Model": Table S2

**Table S2 – Primary Antibodies**

| **Target** | **Species** | **Mono/Poly** | **Company** | **Catalog #** | | **Concentration** |
| --- | --- | --- | --- | --- | --- | --- |
| Laminin β-1 | Rat | Monoclonal (LT3) | Invitrogen | PIMA514657 | | 1:500 |
| Collagen IV | Rabbit | Polyclonal | Abcam | Ab6586 | | 1:500 |
| Perlecan | Rabbit | Polyclonal | Gift from Peter Yurchenco, Rutgers RWJ Medical School | | | 1:1000 |
| E-cadherin | Rat | Monoclonal (DECMA1) | Thermo | | MA1-25160 | 1:1000 |
| Laminin | Rabbit | Polyclonal | Abcam | | Ab11575 | 1:200 (adult sections) |
