## Supplementary material for "A Window into Mammalian Basement Membrane Development: Insights from the *mTurq2-Col4a1* Mouse Model": Table S1

**Table S1 – sgRNA and oligonucleotide sequences**

| **Name** | **Sequence (5’-3’)** |
| --- | --- |
| sgRNA1 for *mTurq2-Col4a1* CRISPR* | GCGCAGCCGAGCAGCTGCGAAG |
| sgRNA2 for *mTurq2-Col4a1* CRISPR | CTGCGAAGGTGAGTTCCCTGCGGG |
| *mTurq2-Col4a1* genotyping F | CCTCCGAGACTGAGCACCTC |
| *mTurq2-Col4a1* genotyping R | TCTCGTTGGGGTCTTTGCTC |
| *mTurq2-Col4a1* genotyping R (het vs hom) | AAATCTCACTGGCTCCTCGGTA |

*sgRNA used to generate the line
