## Supplementary figures and images for "A Window into Mammalian Basement Membrane Development: Insights from the *mTurq2-Col4a1* Mouse Model"

### Supplemental figure 1

Supplemental Figure 1

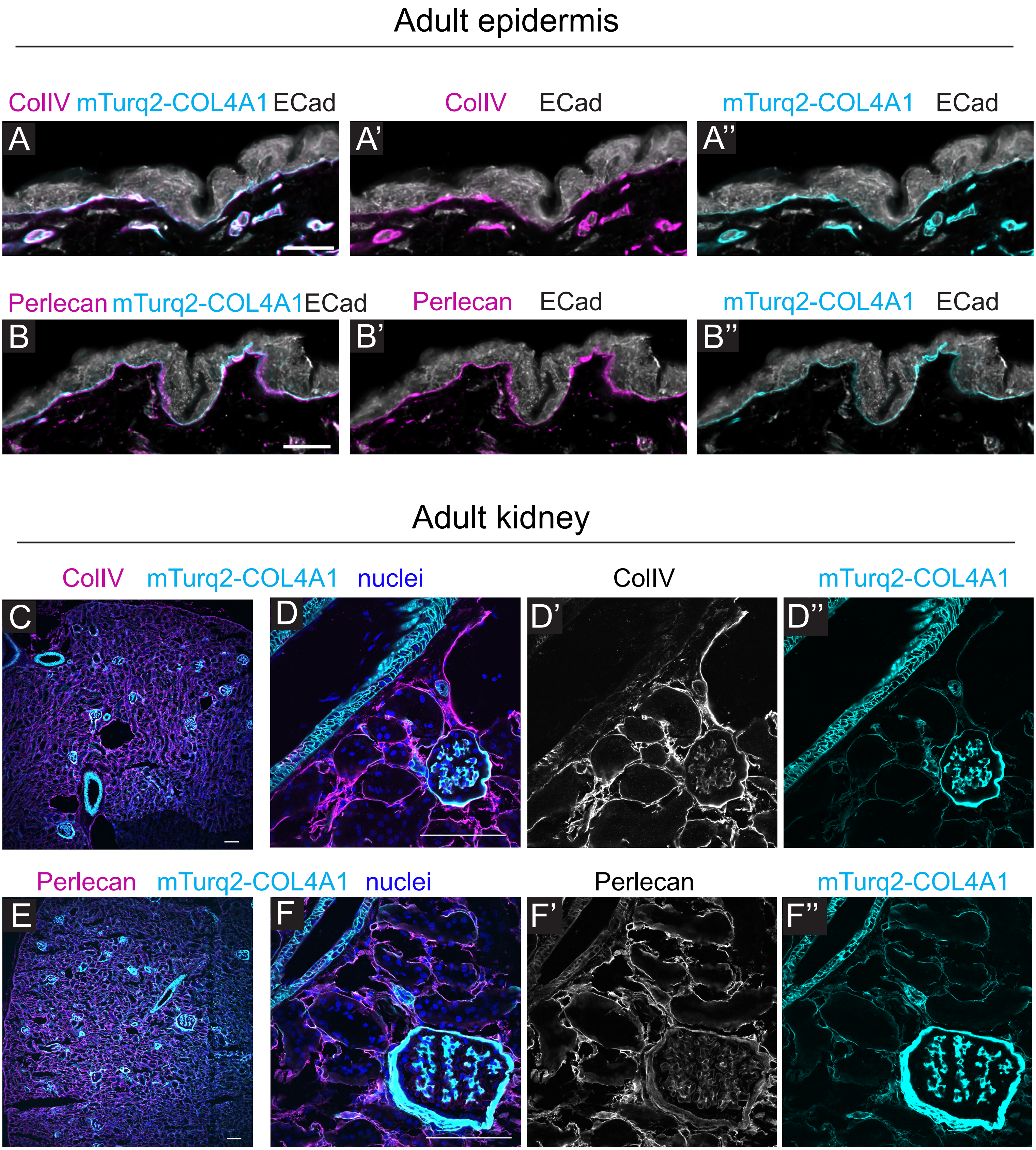

### Supplemental figure 2

#### Supplemental Figure 2

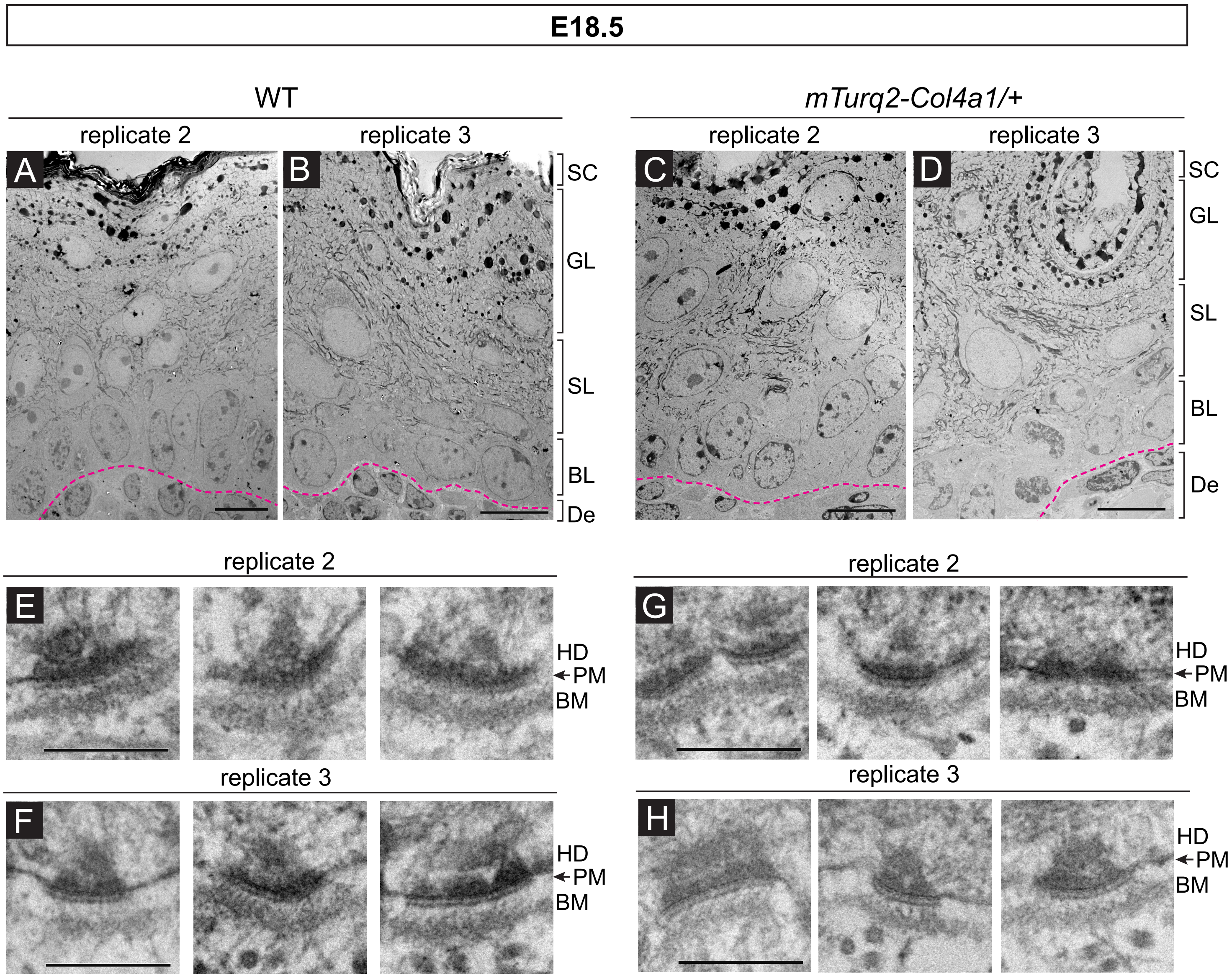

### Supplemental figure 3

#### Supplemental Figure 3

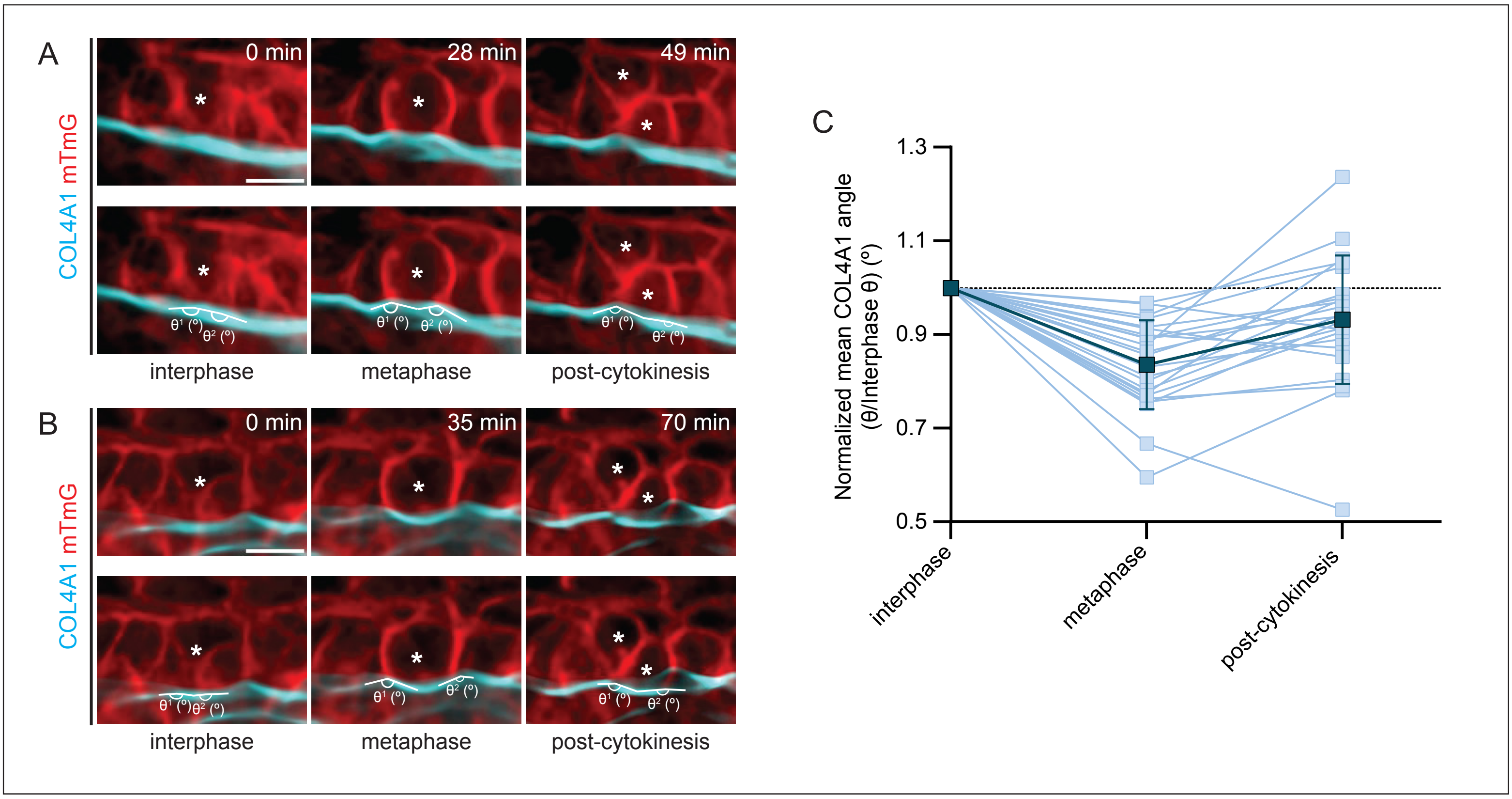
